# AAV-mediated allele-specific silencing alleviates neuropathology in a novel non-human primate model of Spinocerebellar ataxia type 3

**DOI:** 10.1101/2025.10.02.680027

**Authors:** Carina Henriques, Diana Duarte Lobo, Ana Carolina Silva, Ana Rita Fernandes, Miguel Monteiro Lopes, Audrey Fayard, Caroline Jan, Sophie Lecourtois, Martine Guillermier, Julien Flament, João Castelhano, Miguel Castelo-Branco, Rui Caetano Oliveira, Philippe Hantraye, Padmaja Yalamanchili, Steven de Marco, Romina Aron Badin, Luís Pereira de Almeida, Rui Jorge Nobre

## Abstract

Spinocerebellar ataxia type 3 (SCA3), also known as Machado-Joseph disease (MJD), is an autosomal dominant neurodegenerative disorder caused by an abnormal expansion of the cytosine-adenine-guanine (CAG) repeats in the *ATXN3* gene. This mutation results in the production of an Ataxin-3 protein with an extended polyglutamine sequence, contributing to the disease’s neuropathology. Currently, no treatment is available that can slow or halt the progression of SCA3.

Gene-targeted therapies have gained significant attention for their potential to address the root cause of SCA3. Preliminary studies in transgenic mice using adeno-associated viral vector serotype 9 (AAV9) encoding artificial microRNAs targeting the mutant *ATXN3* allele (AAV9-miR-ATXN3) have shown promising results. However, to advance this therapeutic approach toward clinical application, further studies in an animal model that more closely resembles human biology are essential.

In this exploratory study, we assessed the biodistribution and target engagement of AAV9-miR-ATXN3 delivered via intracisterna magna (ICM) injection in non-human primates (NHPs). Using a lentiviral vector (LV) to introduce a mutant Ataxin-3 cDNA with 72 glutamines (LV-mut*ATXN3*-Q72) into the NHP cerebellum, we successfully overexpressed SCA3 in the NHP brain. SCA3 NHP exhibited Ataxin-3 aggregation in the cerebellum, recruitment of inflammatory cells and reduced cerebellar volume. ICM administration of AAV9-miR-ATXN3 effectively directed transgene expression to key brain regions impacted by SCA3 pathology and enabled specific, dose-dependent silencing of mutant Ataxin-3. Furthermore, the therapeutic dose prevented the cerebellar morphological and biochemical alterations induced by the overexpression of mutant *ATXN3*.

These proof-of-concept experiments are crucial, not only for advancing AAV9-miR-ATXN3 toward clinical use but also for establishing a valuable platform for validating future therapeutic interventions.

## Introduction

Spinocerebellar ataxia type 3 (SCA3), commonly referred to as Machado-Joseph disease (MJD), is the most prevalent autosomal dominantly inherited form of ataxia (Bettencourt & Lima, 2011; Schöls et al., 2004). Caused by an unstable repetition of the CAG trinucleotide in exon 10 of the *ATXN3* gene (Costa & Paulson, 2012; Kawaguchi et al., 1994; Rosenberg, 1992; Takiyama et al., 1993), SCA3 is part of the polyglutamine (polyQ) diseases family (La Spada & Taylor, 2003; Orr & Zoghbi, 2007). With an age of onset ranging from 20 to 50 years, SCA3 is an incurable highly debilitating neurodegenerative disorder. Since no therapy is available to cure or delay its progression, treatment is limited to symptomatic and supportive care that seeks to improve patients’ quality of life (Klockgether et al., 2019; H. Paulson & Shakkottai, 1993). Patients typically have a life expectancy of around 20 years after the onset of symptoms, leading to a significant burden of suffering for both patients and their families (Kieling et al., 2007).

During the last few years, different approaches have been explored preclinically to treat SCA3 (reviewed in (Matos et al., 2018)), that range from targeting mutant *ATXN3* nuclei acids (Alves, Hassig, et al., 2008; Alves et al., 2010; Nobre et al., 2022; Nóbrega et al., 2013, 2014; Nóbrega, Codêsso, et al., 2019; Rufino-Ramos et al., 2023), preventing ATXN3 misfolding, aggregation and formation of toxic fragments (Chai et al., 1999; Simões et al., 2012), promoting its proteolytic cleavage or clearance (Fardghassemi et al., 2021; Lee et al., 2010; Lin et al., 2024), restoring dysregulated pathways (Carmona et al., 2017; Cunha-Santos et al., 2016; Duarte-Neves et al., 2015; Koppenol et al., 2023; Nascimento-Ferreira et al., 2011, 2013; Nóbrega et al., 2015; Nóbrega, Mendonça, et al., 2019; Vasconcelos-Ferreira et al., 2022), or fostering neuroprotective mechanisms (Gonçalves et al., 2013).

Nevertheless, owing to its monogenic nature, gene-based approaches that target the mutant *ATXN3* gene or its transcripts remain the most promising (de Sousa-Lourenço et al., 2024). Genome editing approaches, such as zinc finger nucleases (ZFNs), transcription activator-like effector nucleases (TALENs), and clustered regularly interspaced short palindromic repeat (CRISPR)–Cas-associated nucleases, operate at the DNA level, enabling the permanent silencing or correction of the *ATXN3* mutation (Haas et al., 2022; He et al., 2021; Song et al., 2022; Weishäupl et al., 2019). Alternative strategies include ASOs, short chemically modified nucleotide sequences that can either act at the pre-mRNA level, inducing exon skipping or at the mRNA level, resulting in the cleavage of *ATXN3 mRNA* mediated by RNase H (Datson et al., 2017; Evers et al., 2011; Hu et al., 2009, 2011; Kourkouta et al., 2019; McLoughlin et al., 2018, 2023; Moore et al., 2017; Schuster et al., 2024; Toonen et al., 2016). Lastly, RNAi, including short-hairpin RNAs (shRNAs), small interfering RNA (siRNA), and microRNAs (miRNAs), inhibit gene expression by targeting the mutant *ATXN3* mRNA, using the siRNA/miRNA machinery (Alves, Hassig, et al., 2008; Alves et al., 2010; Carmona et al., 2017; Conceição et al., 2016; Martier et al., 2019; Miller et al., 2003; Nobre et al., 2022; Nóbrega et al., 2013, 2014; Nóbrega, Codêsso, et al., 2019; Rufino-Ramos et al., 2023). While editing DNA has made extraordinary progress in recent years, RNA-targeting technology is currently more mature and better suited for *in vivo* gene therapy.

Previously, our group developed an allele-specific RNA interference strategy, specific for an intragenic single-nucleotide polymorphism (SNP) in linkage disequilibrium with the mutant *ATXN3* allele (exon10, rs12895357) (Nobre & Pereira De Almeida, 2020). This strategy has been successfully deployed using an adeno-associated viral vector serotype 9 (AAV9-miR-ATXN3) administered by intraparenchymal, intravenous (Nobre & Pereira De Almeida, 2020) and intracisterna magna (ICM) administration routes (unpublished data) in SCA3 mouse models.

Although the results from mouse models are promising, further studies are required in an animal model closer to humans to scale up and translate this strategy to clinical application. Given the anatomy and functional organisation of the brain, non-human primates (NHP) are the most suitable model to study neurodegenerative disorders, such as SCA3, and for the evaluation of novel therapies (Carlos et al., 2016; Harding, 2013; Kumar & Hedges, 1998; Perelman et al., 2011; Tarantal et al., 2022).

In this work, we conducted a pilot study to model SCA3 in NHP using lentiviral vectors for the delivery and expression of mutant Ataxin-3 cDNA, and to assess the biodistribution and target engagement of AAV9-miR-ATXN3. To the best of our knowledge, this is the first attempt to model SCA3 in *Macaca fascicularis* and to evaluate a therapeutic approach in a model phylogenetically closer to humans. This understanding is crucial, not just to aid the clinical translation of AAV9-miR-*ATXN3*, but also, as a platform for the validation of other novel therapies and biomarkers.

## Methods

### In vitro experiments

#### Lentiviral vector production

Lentiviral vectors (LVs) were generated using a previously cloned cDNA encoding the human mutant Ataxin-3, isoform mjd1a, with 72 glutamines (Q) (mut*ATXN3*-Q72), in the transfer vector self-inactivating (SIN), carrying the phosphoglycerate kinase (PGK) promoter (SIN-W-PGK) (Alves, Régulier, et al., 2008).

Human embryonic kidney 293 cells stably expressing the SV40 large T antigen (HEK 293T) were used for LVs production using the four-plasmid protocol previously described (De Almeida et al., 2002). Succinctly, 24 hours before transfection, 1.05 x 10^7^ HEK 293T cells were plated into 15-cm dishes and cultured in Dulbecco’s Modified Eagle Medium (DMEM) high glucose (Thermo Fisher Scientific) supplemented with 10 % fetal bovine serum (FBS, Biowest), 1 % Penicillin/Streptomycin (P/S, Life Technologies). Cells were transfected with pCMVDR-8.92, pMD.G, pRSV-Rev, and SIN-W-PGK transfer vectors using polyethylenimine (PEI) linear MW 40000 (Polysciences). Forty-eight hours later, the cell-conditioned medium was filtered with 0.45 µm polyvinylidene fluoride (PVDF) filters (Merck Millipore) and concentrated by two ultracentrifugation steps at 70 000 g (Optima XE-100, Beckman Coulter). LVs were resuspended in phosphate-buffered saline (PBS, Fisher BioReagents) with 1% bovine serum albumin (BSA, Merck Millipore). LV concentration was determined by P24 antigen enzyme-linked immunosorbent assay (RETRO-TEK HIV-1 P24 ELISA kit, ZeptoMetrix), following the manufacturer’s instructions. Concentrated viral stocks were stored at-80 °C until use.

#### Adeno-associated viral vector production

The therapy involves the use of an AAV9 containing a self-complementary expression cassette with two copies of previously validated artificial microRNAs that specifically target the mutant *ATXN3* allele (Nobre & Pereira De Almeida, 2020), under the control of the cytomegalovirus enhancer/chicken beta-actin (CAG) promoter (AAV9-miRNA-ATXN3). AAV9 vectors were produced by PackGene Biotech, LLC (17 Briden Street, Rm 309, Worcester, MA 01605, USA). Concentrated viral stocks were suspended in sterile PBS/0.001% Pluronic F-68 (F68) solution.

AAV viral genomes (vg) per µL were assessed by detecting the copy numbers of AAV2 ITRs by Real-time SYBR Green PCR. Viral stocks were stored at-80 °C until use.

### In vivo Experiments

#### Animals

Twelve 9-year-old, wild-type, naïve adult male cynomolgus monkeys (*Macaca fascicularis*, supplied by Camarney, Spain) were recruited in the study. The animals were housed under standard conditions, on a 12-hour light/dark cycle, in a temperature and humidity-controlled room (22 ± 1 °C and 50 %, respectively). Water and food were available *ad libitum*. All animal studies were conducted according to European regulations (EU Directive 2010/63/EU) and in compliance with Standards for Humane Care and Use of Laboratory Animals of the Office of Laboratory Animal Welfare (OLAW—no. #A5826–01) in a facility authorised by local authorities (authorisation no. #D92-032-02). The experimental protocol was reviewed and approved (authorisation no. A21_019) by the local ethics committee (CETEA No.44). A detailed weekly physical examination, including body weight evaluation, was performed to monitor general health. All efforts were made to minimise animal distress or suffering. Animal care was guaranteed by veterinarians and animal technicians proficient in the healthcare and housing of NHPs.

#### In vivo injection of viral vectors

For the AAV9-miR-ATXN3 biodistribution study, six cynomolgus monkeys were randomised into two groups of three animals. Two doses of AAV9-miR-ATXN3, 0.3 x 10^13^ vg (d1) or 3 x 10^13^ vg (d2), were delivered via ICM injections (Figure 1.a and Supplementary Table 1).

**Figure 1.**
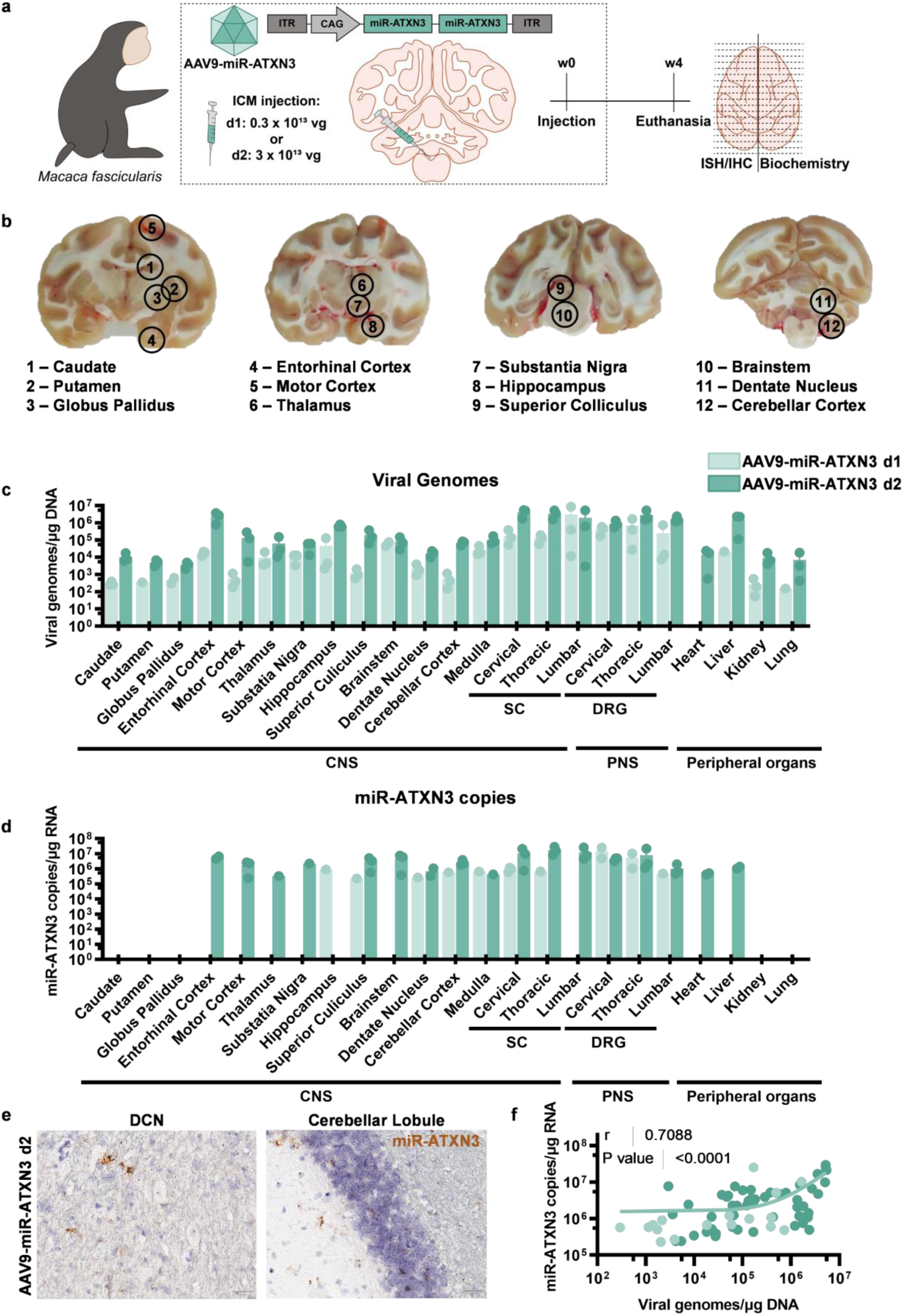
I**n**tracisterna **magna administration of AAV9-miR-ATXN3 led to transgene expression in crucial areas of SCA3 pathology. (a)** Schematic representation of the experimental procedure. Six eight-year-old wild-type male cynomolgus monkeys were injected with 3 x 10^12^ vg (d1, n = 3) or 3 x 10^13^ vg (d2, n = 3) of AAV9-miR-ATXN3. Four weeks post-injection monkeys were euthanised for posterior immunostaining, *in situ* hybridization and biochemical analysis. **(b)** Representative images of NHP brain slices and the respective localization of brain punches used for biochemical analysis. **(c)** AAV viral genomes in different regions, relative to µg of TOTAL DNA. Results are shown as mean ± SD for AAV9-miR-ATXN3 d1 and AAV9-miR-ATXN3 d2 (n = 1-3, per group). **(d)** Artificial miRNA copies in different regions, relative to µg of TOTAL RNA. Results are shown as mean ± SD for AAV9-miR-ATXN3 d1 and AAV9-miR-ATXN3 d2 (n = 1-3, per group). **(e)** Monkey coronal brain sections were incubated with nucleotide probes specific for miR-ATXN3. Scale bar = 100 µm. **(f)** Pearson’s r correlation analysis between viral genomes and miR-ATXN3 copies, considering all the different dissected regions from the two test groups. Data points are shown for AAV9-miR-ATXN3 d1 (n= 17) and AAV9-miR-ATXN3 d2 (n = 53). CNS = central nervous system; DCN = deep cerebellar nuclei; DRG = dorsal root ganglions; PNS = peripheral nervous system; SC = spinal cord.

To develop a viral-based NHP model of SCA3 and simultaneously evaluate AAV9-miR-ATXN3 target engagement, six additional cynomolgus monkeys were randomized into three groups of two animals. NHP received cerebellar intraparenchymal (IP) administration of a single dose of LV-mut*ATXN3* (20 000 ng of P24), or LV buffer (PBS/BSA 1%), and a single ICM injection of 2 x 10^13^ vg of AAV9-miR-ATXN3, or AAV buffer (PBS/0.001% F68 solution) (Figure 2.a and Supplementary Table 2).

**Figure 2.**
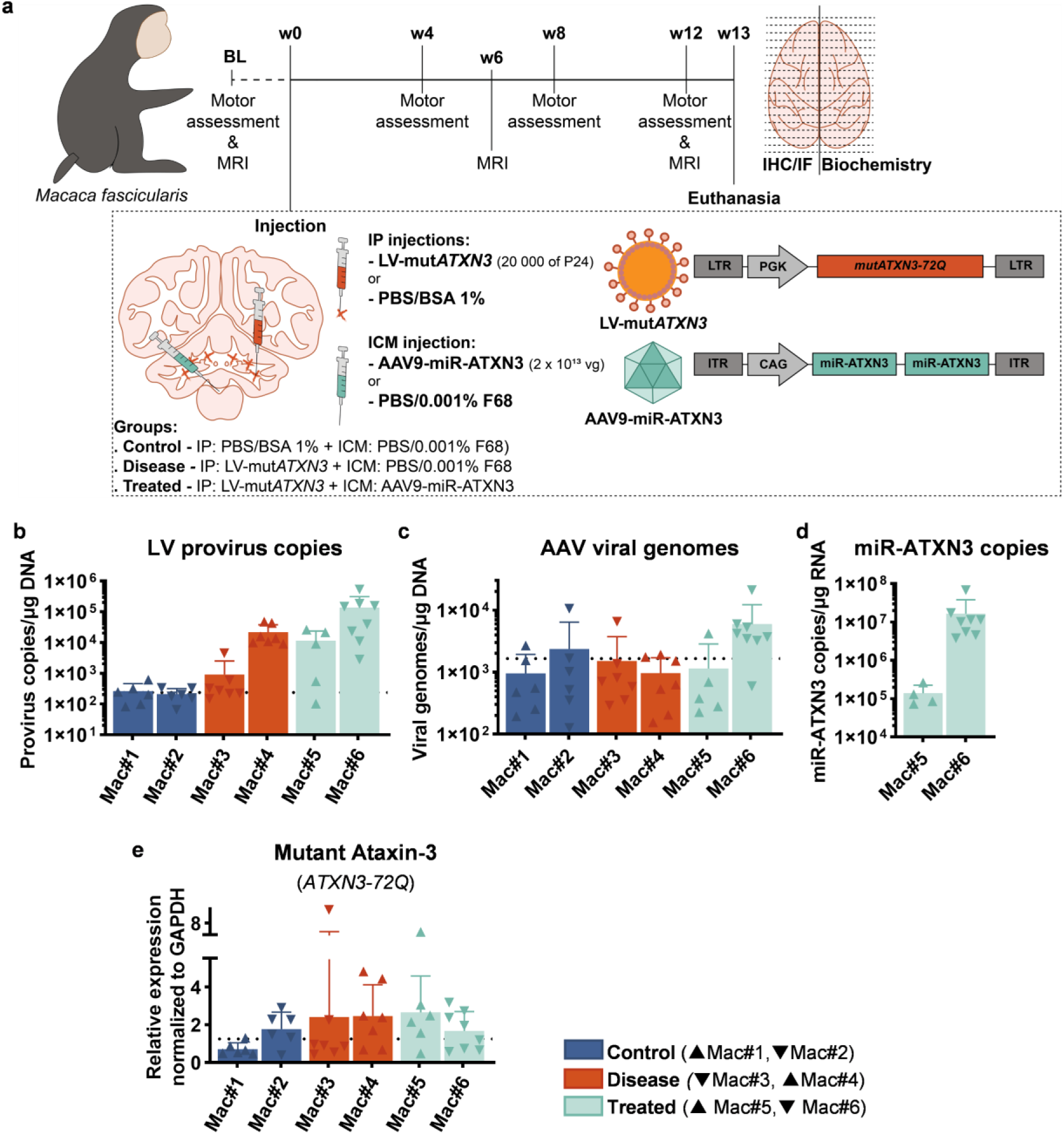
A**A**V9**-miR-ATXN3 can specifically silence mut*ATXN3* overexpressed in the cerebellum of WT NHP in a dose-dependent manner. (a)** Schematic representation of the experimental procedure. Six ten-year-old wild-type male cynomolgus monkeys were divided into three study groups: Control, disease and treated. Monkeys in the treated group (Mac#5 and Mac#6) received six injections of LV encoding for human mutant Ataxin-3 with 72 glutamines (Q) (LV-mut*ATXN3*) in the cerebellum and an intracisterna magna (ICM) injection of AAV9-miR-ATXN3. Monkeys in the disease group (Mac#3 and Mac#4) received six injections of LV-mut*ATXN3* in the cerebellum and an ICM injection of PBS/0.001% Pluronic F68, AAV’s buffer, and NHP in the third group, Ctrl (Mac#1 and Mac#2), received six injections of PBS/BSA 1 % in the cerebellum and an ICM injection of PBS/0.001% Pluronic F68. Motor behaviour was evaluated, and brains were imaged by magnetic resonance imaging (MRI) in the baseline (BL) and throughout the *in vivo* phase (every four weeks and every six weeks, respectively). Thirteen weeks post-injection monkeys were euthanised for posterior immunostaining and biochemical analysis. For biochemical analysis, six to eight small pieces were obtained from the right cerebellum of each animal, posteriorly used for qPCR (**b, c)**, and RT-PCR (**d, e)**. (**b, c**) LV provirus copies and AAV viral genomes per µg of DNA, respectively. (**d**) RT-PCR analysis of miR-ATXN3 copies relative to µg of RNA**. (e)** Expression levels of mutant Ataxin-3 (*Macaca fascicularis*) (*ATXN3*) mRNA, relative to *Macaca fascicularis GAPDH*. Results are shown as mean ± SD for (b,c,d,e) Control, mut*ATXN3*, and mut*ATXN3* + AAV9-miR-ATXN3 (n = 5-8, per animal). The dotted line represents the average of the control group.

The animals were induced with intramuscular (IM) injections of ketamine (10 mg/kg) and xylazine (1 mg/kg) and maintained anaesthetised with an intravenous (IV) administration of propofol (1 ml/kg/h). Temperature, respiratory frequency, exhaled CO_2_, heart rate and blood pressure were constantly monitored during the surgical procedure. Buprenorphine (0.02 mg/kg) and a wide-spectrum antibiotic (Clamoxyl long-acting, 20 mg/kg) were administered immediately before and after surgery. Since Mac#4 had an allergic reaction to the anaesthetics, this animal was treated with atropine (0.02 mg/kg) and induced with a combination of ketamine (1 mg/kg) and medetomidine (0.5 mg/kg).

Magnetic resonance imaging (MRI) anatomical scans (described below) were performed to determine IP injection coordinates. IP administration was performed bilaterally, in a total of six injection sites (three per hemisphere), in the white matter (WM) near the dentate cerebellar nuclei (DCN). Precise injection coordinates were obtained by employing the frameless stereotaxic neuronavigation system Brainsight (Rogue Research Inc) (Wetzel et al., 2019) and injections were performed using a Hamilton syringe connected to a KDS injection micropump. At each coordinate, a single deposit of 15 μL was injected at a rate of 3 μL/min.

For ICM administration, animals were placed in a stereotaxic frame with the nose pointing down to expose the cervical area. The occipital protuberance and the transverse process of the atlas (C1) were palpated to determine the insertion point. A Quincke spinal needle with 22 G and 1–1.5-inch (Becton Dickinson) fixed onto a stereotaxic micromanipulator was advanced into the suboccipital space at an angle of 30°. Needle placement was confirmed by cerebrospinal fluid (CSF) return and 1.5 mL of CSF was collected. Then a syringe was attached to the needle through a 3-way valve. Injection was performed slowly by hand over one minute and the linker was flushed with 200 μL of the primate’s own CSF.

#### Magnetic resonance imaging

MRI was performed before surgery (baseline, BL), and it was repeated at 6 and 12 weeks post-injection (p.i.) in the 6 animals used for AAV9-miR-ATXN3 target engagement evaluation. All MRI images were acquired on a 7 Tesla Bruker scanner with a 13.5 cm surface coil placed over the top of the head for radiofrequency transmission and reception. A high isotropic resolution image was acquired using FLASH sequence (TE/TR = 5/2315 ms, 8 averages, in-plane resolution = 75 x 75 μm², 140 slices with 75 µm thickness, Tacq = 45 min) for skull reconstruction. These images were co-registered with anatomical T2-weighted images (Multi Slices Multi Echoes sequence (TE/TR = 10/3500 ms, 5 echoes, effective TE = 30 ms, in-plane resolution = 500 x 500 μm², 60 slices with thickness = 1 mm, Tacq = 15 min)) for neuronavigation.

#### Cerebellar volume analysis

T2 MRI images were processed using SMP12 (version 7771), Mango (version 4.1) and PMOD (version 4.205) software. Initially, T2 DICOM images were converted to NIFTI-1 format using the DICOM import function in SPM12. The images were then reoriented to the desired anatomical alignment and skull stripping was performed using the Brain Extraction Tool (BET) plugin in Mango.

In the absence of a T2-weighted cerebellar template for *Macaca fascicularis*, the INIA19 whole-brain template for *Macaca mulatta* (Rohlfing et al., 2012) was used. Using the PMOD Neuro Tool, regions corresponding to the right and left cerebellum in the INIA19 template were merged to define the right and left cerebellar volumes of interest (VOIs), respectively.

Next, the PMOD Fusion Tool was used to register the INIA19 template to itself, followed by the transformation of the co-registered template to match the skull-stripped images using the Deform method. The same transformation was applied to the cerebellar VOIs, previously created with the PMOD Neuro Tool. Cerebellar volumes were calculated by co-registering the transformed cerebellar VOIs with the transformed INIA19 template.

The right and left cerebellar volumes were combined to calculate the total cerebellar volume. To account for interindividual neuroanatomical differences, total cerebellar volume was normalised as a percentage of baseline (BL). Default parameters were used in the various operations.

#### Behavioural Assessment

To evaluate macaque locomotor activity in the target engagement study (n = 6), 30-minute videos were acquired in a dedicated room equipped with a transparent glass cage. Video recordings were performed at BL and every four weeks after stereotaxic injection (4, 8, and 12 weeks p.i.). Videos were analysed using Ethovision software (Noldus). To discard intraindividual variability and stereotypic behaviours, total distance travelled, and mean velocity are expressed as a percentage of BL. The mean of five videos was used to calculate BL locomotion parameters.

#### Tissue Collection and Preparation

Four or thirteen weeks p.i. macaques were anaesthetized with ketamine (10 mg/kg) and xylazine (0.05 mg/kg) and administered a dose of buprenorphine (0.02 mg/kg) for analgesia before termination by pentobarbital overdose (180 mg/kg, IV), and transcardial perfusion with 0.9% NaCl. After death, quickly excised brains were divided into two halves and sliced into 4 mm coronal slabs. The right hemisphere was assigned for biochemistry and the left hemisphere for histological staining, immunostaining, and *in situ* hybridization (ISH).

In the right hemisphere of animals from the AAV9-miR-ATXN3 biodistribution assay, thirteen brain regions (caudate, putamen, globus pallidus, entorhinal cortex, motor cortex, thalamus, substantia nigra, hippocampus, superior colliculus, brainstem, dentate nucleus, cerebellar cortex and medulla) were punched, flash-frozen and stored at-80 °C. Cervical, thoracic and lumbar dorsal root ganglions (DRG) and spinal cord (SC), as well as heart, lung, kidney and liver punches, were also collected for biochemistry analysis. Samples were sent to Charles River Labs for posterior analysis (Reno, NV USA).

For the generation of the NHP SCA3 viral-based model and AAV9-miR-ATXN3 target engagement experiment, right hemisphere slabs were stored at-80 °C, and six to eight cerebellar samples were excised in dry ice for biochemical analysis.

In both experiments, the left hemisphere was post-fixed with 4 % paraformaldehyde (PFA) at 4 °C for one week. For histological staining and immunostaining, left hemisphere 4 mm slices were transferred to sucrose gradients in PBS for cryoprotection for at least 48 hours. Flash-frozen in dry ice, coronal sections were subsequently cut at 40 μm thickness, using a cryostat (CryoStar NX50, Thermo Scientific) at-20 °C. Sections were stored in 15-well trays as free-floating sections in 30% glycerol, 30% ethylene glycol and 10% phosphate buffer solution. Trays were stored at-20 °C until further processing. For ISH, left hemisphere 4 mm slices were transferred to PBS. The miRNAscope ISH assay, with customised probes for miR-ATXN3, was performed by Advanced Cell Diagnostics (7707 Gateway Blvd, Newark, CA 94560, USA).

#### Immunostaining

For immunohistochemistry (IHC), free-floating sections were rinsed in PBS and incubated in 0.3 % hydrogen peroxide (Fisher Chemical) in distilled water for 20 min to block endogenous peroxidases. Tissue blocking and permeabilization were subsequently performed in a blocking solution (0.2 % Triton X-100, Sigma-Aldrich) containing 4.5 % Normal Goat Serum (NGS, Gibco) in PBS), for 20 min at RT. Brain slices were then incubated for two overnights at 4 °C with primary antibody mouse anti-Ataxin-3, 1h9 (1:1 000 Millipore, MAB5360) diluted in 0.2 % Triton X-100 and 3 % NGS in PBS. Free-floating sections were washed in PBS and incubated for 1 h at RT with the respective biotinylated secondary antibody diluted in 0.2 % Triton X-100 and 3 % NGS in PBS (goat anti-mouse, 1:250, Vector Laboratories, BA-9200). Subsequently, bound antibodies were visualised in rinsed free-floating sections by the Avidin-Biotin complex (ABC) amplification system (Vectastain ABC kit, Vector Laboratories) using 3,3′-diaminobenzidine tetrahydrochloride (DAB Substrate Kit, Vector Laboratories) as substrate. The reaction was stopped by washing sections in PBS, after achieving optimal staining. Sections were mounted on gelatine-coated slides, and dehydrated in an ascending ethanol sequence (50 %, 70 %, 95 % and 100 %, Thermo Scientific). Slides were then cleared with xylene solution (Thermo Scientific) and finally coverslipped with Richard-Allan Scientific Mounting Medium (Thermo Scientific).

For immunofluorescence (IF), free-floating sections were rinsed in PBS and incubated in a blocking solution (0.2 % Triton X-100 containing 4.5 % NGS) in PBS) for 30 min at room temperature (RT) for blocking and permeabilization. After blocking, brain slices were incubated overnight at 4 °C with primary antibody mouse anti-Ataxin-3, 1h9 (1:1 000 Millipore, MAB5360) diluted in 0.2 % Triton X-100 and 3 % NGS in PBS. Following three washing steps in PBS, free-floating sections were incubated for 1 hour at RT with the respective secondary antibody diluted in 3 % NGS in PBS (568 goat anti-mouse, 1:250, Invitrogen, A11004). Subsequently, free-floating sections were rinsed and stained with 4′,6-diamidino-2-phenylindole (DAPI, Thermo Scientific) for 8 min. Free-floating sections were rinsed and mounted on gelatine-coated slides with fluorescence mounting medium (DAKO).

Images were acquired using a Zeiss Axio Scan.Z1 microscope, equipped with a Plan-Apochromat 20X/0.8 air objective.

#### Cresyl violet staining

For cresyl violet staining, after being rinsed in PBS, free-floating sections were mounted on gelatine-coated slides and dried at RT. To stain the Nissl substance present in the neuronal bodies, sections were immersed in cresyl violet solution for 5 min. Then, slides were differentiated in 2 % acetic acid following an ascending ethanol sequence (50 %, 70 %, 95 % and 100 %, Thermo Scientific). After a clearing step in xylene (Thermo Scientific), sections were mounted with Richard-Allan Scientific Mounting Medium (Thermo Scientific).

Images were acquired using a Zeiss Axio Scan.Z1 microscope, equipped with a Plan-Apochromat 20X/0.8 air objective.

#### DNA, RNA and protein extraction

Simultaneous DNA, RNA and protein extractions were performed with AllPrep DNA/RNA/Protein Mini Kit (Qiagen), according to the manufacturer’s instructions. Before extraction, tissues were homogenised with a syringe with a 21 G needle (BRAUN) in lysis buffer. DNA and RNA were eluted in 40 µL of DNase/RNase-free water and protein extracts were dissolved in 8 M Urea (Fisher Chemical) in 100 mM Tris-HCl (Fisher BioReagents), at pH = 8, supplemented with 1% sodium dodecyl sulfate (SDS, Fisher BioReagents) and protease inhibitor cocktail (complete Mini, Roche Diagnostics). RNA and protein extracts were stored at-80 °C and DNA was stored at-20 °C.

Before use, purified DNA and RNA were quantified by optical density (OD) using a Nanodrop 2000 Spectrophotometer (ThermoFischer Scientific) and protein concentration was determined, after two cycles of sonication at 20-40 kHZ with 8-12 pulses of 1-3s, by bicinchoninic acid (BCA) according to the manufacturer’s instructions (Bio-Rad Laboratories).

#### AAV genome copy number quantification

The AAV genome copy number was determined using the AAVpro Titration Kit (Takara Bio) according to the manufacturer’s instructions. Briefly, genomic DNA was diluted to 10 ng/μL and used as a template for real-time qPCR. Each reaction contained a total of 50 ng of DNA and was performed alongside a standard curve generated from serial dilutions of a calibrated AAV control template. The ITR sequence of AAV2 was targeted to quantify the viral genome in each sample. Sample Ct values were correlated to the standard curve to determine the AAV genome copy number. Viral genome values were then normalised to total DNA (vg/μg).

#### Lentiviral provirus quantification

The lentiviral provirus copy number per diploid cell was determined using the Lenti-X Provirus Quantitation Kit (Takara Bio), according to the manufacturer’s instructions. Briefly, DNA was diluted to 10 ng/µL and used as a template for real-time qPCR. Each reaction contained a total of 20 ng of total DNA and was performed alongside a standard curve generated from serial dilutions of a calibrated provirus control template. To determine provirus copy, raw Ct values of the samples were correlated to the standard curve to determine the provirus copy number. A correction coefficient due to different amplification efficiencies of genomic DNA and plasmids was applied and the number of total cells was extrapolated from the total amount of DNA, as recommended by the manufacturers.

#### cDNA synthesis and RT-PCR

For mRNA quantification experiments, 125-250 ng of RNA was first treated with DNase (Qiagen). Next, RNA samples were reverse transcribed into double-stranded cDNA using the iScript cDNA Synthesis kit (Bio-Rad Laboratories), according to the manufacturer’s recommendations. cDNA was stored at 4 °C if used within 2 days and at-20 °C for long-term storage.

Real-time PCR was carried out in a CFX96 Touch Real-Time PCR Detection System (Bio-Rad Laboratories), using 96-well microliter plates and the SsoAdvanced Universal SYBR Green Supermix (SsoAdv, Bio-Rad Laboratories), following the manufacturing protocol. Briefly, real-time PCR was carried out in 10 µL reaction volume, with 2.5 ng / µL of cDNA, 0.5x SsoAdv and 10 µM of forward and 10 µM of reverse primers (Supplementary Table 3). All reactions were performed in duplicate: 95 °C for 30 s, followed by 40 cycles at 95 °C for 5 s and 15 s at the corresponding annealing temperature. The melting curve was performed with slow heating, starting at 65 °C with an increment of 0.5 °C per 5 s up to 95 °C. *GAPDH* was used as a reference gene. The mRNA change to control samples was determined by the 2^delta-delta Ct method.

For miRNA quantification experiments, specific cDNAs were synthesised using a TaqMan MicroRNA Reverse Transcription Kit combined with specific TaqMan MicroRNA Assays (Applied Biosystems, Thermo Fisher Scientific) for each miRNA, according to the manufacturer’s instructions. Briefly, 5 ng of RNA was used for reverse transcription with primers specific for miR-ATXN3 (assay ID CTKA3RH). Real-time qPCR was performed using TaqMan Fast Advanced Master Mix (Applied Biosystems) in a StepOnePlus Real-Time PCR System (Applied Biosystems). For the determination of miRNA absolute expression levels, a standard curve was performed using a commercially synthesised RNA oligonucleotide (Thermo Scientific). Finally, microRNA levels were normalised to microRNA copies/μg of total RNA.

#### Western blot

For protein analysis, 40 µg of protein extracts were resolved in SDS-polyacrylamide gels (4 % stacking and 10 % running) and transferred onto PVDF membranes (Immobilon-P, Merck Millipore), according to standard protocols. After protein electrotransfer, membranes were stained with No Stain (Thermo Fisher Scientific) to visualise total protein. Then, membranes were blocked by incubation in 5 % non-fat milk powder in 0.1 % Tween 20 (Fisher BioReagents) in Tris buffered saline (TBS-T), for 1 h at RT. Immunoblotting was performed overnight at 4 °C with the primary antibody mouse anti-Ataxin-3, 1h9 (1:1 000 Millipore, MAB5360), followed by RT incubation with corresponding alkaline phosphatase-coupled secondary antibody (goat anti-mouse, 1:10 000, Invitrogen, 31328). The presence of the antigens of interest was observed with Enhanced Chemifluorescence substrate (ECF, Ge Healthcare) and chemifluorescence imaging (ChemiDoc MP imaging system, Bio-Rad Laboratories).

## Statistical Analysis

All statistical analyses were performed using the GraphPad Prism software (version 9.0.0). Data are presented as mean ± standard deviation (SD). Simple linear regression and Pearson’s correlation were conducted where applicable. Statistical significance was determined using the following thresholds: * *p* < 0.05, ** *p* < 0.01, *** *p* < 0.001, and **** *p* < 0.0001.

## Results

### Intracisterna magna administration of AAV9-miR-ATXN3 effectively induces transgene expression in the crucial brain areas of SCA3 pathology

Firstly, we aimed to assess the biodistribution of AAV9 carrying two copies of an allele-specific miRNA expression cassette targeting a SNP in linkage disequilibrium with the mutant *ATXN3* allele (exon10, rs12895357 SNP) (AAV9-miR-ATXN3) after ICM delivery in NHP. For that, six *Macaca fascicularis* were randomised to receive a single dose of AAV9-miR-ATXN3 at either 0.3 x 10^13^ vg (AAV9-miR-ATXN3 d1) or 3 x 10^13^ (AAV9-miR-ATXN3 d2) viral genomes (vg) per animal (Figure 1.a). Four weeks after vector administration, animals were euthanised, and samples were collected throughout the brain (Figure 1.b), spinal cord (SC), and dorsal root ganglions (DRG), as well as peripheral organs, including the heart, liver and kidneys. AAV transduction efficiency was assessed by evaluating AAV viral genomes (Figure 1.c) and miR-ATXN3 copy number (Figure 1.d).

For both high (d2) and low (d1) AAV doses, the highest number of viral genomes per genomic DNA was observed in the SC and DRG (Figure 1.c). Conversely, the lowest levels of viral genomes were observed in the basal ganglia, namely in globus pallidus, putamen and caudate, and in the lungs and kidneys. For animals injected with the high dose (d2), the entorhinal cortex and liver also exhibited high levels of AAV transduction.

Regarding miR-ATXN3 copy number, for monkeys that received the low dose of AAV9-miR-ATXN3 (d1), the highest levels were observed in the cervical SC and the cervical and thoracic DRG (Figure 1.c). Samples from cervical, thoracic and lumbar SC were the most enriched in miR-ATXN3 in animals from the high dose (d2) group. Moreover, in both groups, it was possible to detect miR-ATXN3 in the dentate nucleus and cerebellar cortex. On the other hand, in peripheral organs, miR-ATXN3 expression was only detectable in monkeys which received the higher AAV dose. The expression of miR-ATXN3 in the cerebellum (DCNs and lobules) was further confirmed by *in situ* hybridisation (ISH) (Figure 1.e).

Considering all the different dissected regions from the two test groups (n = 70), there was a strong correlation between viral genomes and miR-ATXN3 transgene expression, as indicated by Pearson’s r correlation analysis (r = 0.7088, *p* < 0.0001) (Figure 1.f). Moreover, a simple linear regression analysis showed that the model explained 50.24% of the variance (r^2^ = 0.5024) (Supplementary Table 4).

No differences were observed in glial fibrillary acidic protein (GFAP) immunoreactivity in different brain regions between animals treated with high (d2) and low (d1) doses of AAV9-miR-ATXN3 (Supplementary Figure 1).

Overall, these results indicate that AAV9-miR-ATXN3 delivered by ICM injection is safe and efficiently transduces the most affected brain regions in SCA3, namely the dentate nucleus, cerebellar cortex, spinal cord and brainstem without causing significant neuroinflammation.

### AAV9-miR-ATXN3 specifically silences mutATXN3 overexpressed in the cerebellum of NHP in a dose-dependent manner

After validating the transduction of the affected areas in SCA3 through injection into the *cisterna magna*, particularly with the highest dose, we conducted a second experiment to evaluate the target engagement of miR-ATXN3 in an NHP SCA3 LV-based model. To achieve this, six ten-year-old male *Macaca fascicularis* were randomised into three study groups: (1) “Control group”, received LV’s buffer into the cerebellum and AAV’s buffer by ICM administration (Mac#1 and Mac#2); (2) “Disease group” received a single dose of LV-*mutATXN3* into the cerebellum and AAV’s buffer by ICM administration (Mac#3 and Mac#4); (3) “Treated group”, received a single dose of LV-mut*ATXN3* into the cerebellum and an ICM injection of AAV9-miR-ATXN3 (Mac#5 and Mac#6) (Figure 2.a). Before injection, NHP underwent motor behaviour assessment and anatomical brain imaging. After viral administration, motor behaviour was assessed every four weeks and magnetic resonance imaging (MRI) was performed every six weeks. Moreover, a detailed weekly physical examination was carried out to assess animal welfare. Animals were euthanised at 13 weeks post-injection for posterior biochemical and immunostaining analysis.

Firstly, we evaluated LV and AAV transduction, by assessing LV provirus copies (Figure 2.b) and AAV viral genomes (Figure 2.c). A high variability was observed in LV provirus copies among the different animals, even within the same study group (Figure 2.b). For instance, in the disease group, Mac#4 had 24 times more LV provirus copies per μg of DNA when compared to Mac#3 (21 553 ± 16 622 and 908.3 ± 1 615, respectively). In the treated group, Mac#6 had 12 times more LV provirus copies per μg of DNA when compared to Mac#5 (138 311 ± 177 773 and 11 584 ± 12 071, respectively).

High variability in AAV viral genomes was also observed in the treated group, with the mean number per μg of DNA from different cerebellar regions of Mac#5 falling below the detection limit (Figure 2.c). Despite that, it was possible to detect miR-ATXN3 in brain samples from both monkeys in this group (Mac#5 and Mac#6) (Figure 2.d), with no amplification observed in the control and disease groups. However, there was insufficient evidence to suggest a correlation or a relationship between viral genomes and miR-ATXN3 transgene expression, as indicated by Pearson’s r (r = 0.1708, p = 0.6155) and r² (r² = 0.02919) (Supplementary Table 4).

In summary, based on LV provirus copies per μg of DNA, the injected animals can be classified into three profiles: low (Mac#3), medium (Mac#4 and Mac#5), and high (Mac#6) dose. A similar pattern was observed in miR-ATXN3 levels, with Mac#5 exhibiting a low number of miR-ATXN3 copies per μg of RNA, while Mac#6 had a substantially higher amount. Next, we evaluated mut*ATXN3* mRNA expression by RT-qPCR (Figure 2.e). In the treated group, the animal with the lowest number of AAV viral genomes and miR-ATXN3 copies (Mac#5) had levels of mut*ATXN3* mRNA similar to animals in the disease group. In contrast, the treated animal with the highest number of AAV viral genomes (Mac#6) showed a reduction in mut*ATXN3* expression, reaching residual levels near the detection limit, comparable to those observed in the control group.

Unfortunately, the levels of mutant Ataxin-3 protein and Ataxin-3 protein aggregates in the cerebellar homogenates were too low to be clearly detected by Western blot (Supplementary Figure 2.a). We also evaluated whether the overexpression of mut*ATXN3* would impact the levels of endogenous Ataxin-3. No significant differences were observed in endogenous *Macaca fascicularis* Ataxin-3 mRNA or protein levels (Supplementary Figure 2).

Then, we investigated if Ataxin-3 inclusions could be found in the cerebellum of NHP injected with LV-mut*ATXN3*. For that, we performed IHC and IF for Ataxin-3 antibody 1h9 in cerebellar coronal sections (Figure 3.a, b). Ataxin-3 aggregates were observed in the animal with the highest dose of LV from the disease group (Mac#4) and in the animal with the lowest levels of miR-ATXN3 from the treated group (Mac#5) (Figure 3.a, b). In contrast, in the animal with the highest levels of miR-ATXN3 (Mac#6), no inclusions were detected (Figure 3.a, b). Moreover, the presence of Ataxin-3 aggregates was associated with an inflammatory response, as revealed by cresyl violet staining (Figure 3.a, c).

**Figure 3.**
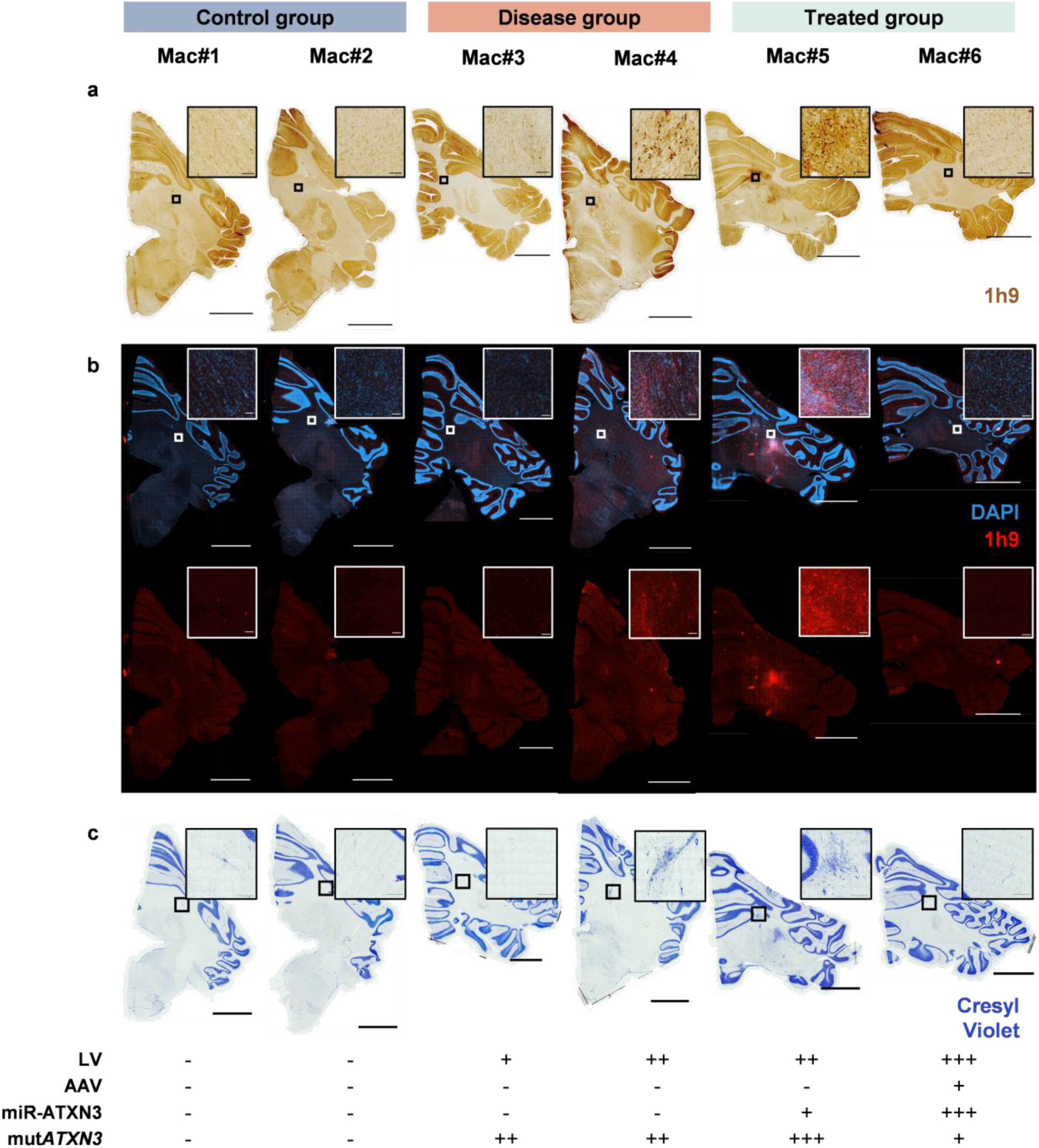
A**t**axin**-3 aggregation in SCA3 NHP and recruitment of inflammatory cells to the affected regions.** Coronal cerebellar sections of monkeys from control, disease, and treated groups were incubated with Ataxin-3 antibody 1h9 for immunofluorescence **(a)** and immunohistochemistry **(b)** and stained with cresyl violet **(c)**. **(b)** Anti-1h9, red; DAPI, blue. **(a-b)** Scale bar = 5000µm; 100 µm; **(c)** Scale bar = 5000µm; 100µm.

Overall, these results suggest that AAV9-miR-ATXN3 can silence mut*ATXN3* in a dose-dependent manner.

### Phenotypic heterogeneity among NHP overexpressing mutant Ataxin-3 in the cerebellum

Given that the primary clinical features of SCA3 are linked to the loss of motor functions, locomotor activity was assessed at baseline and every four weeks after injection (Figure 4.a - c).

**Figure 4.**
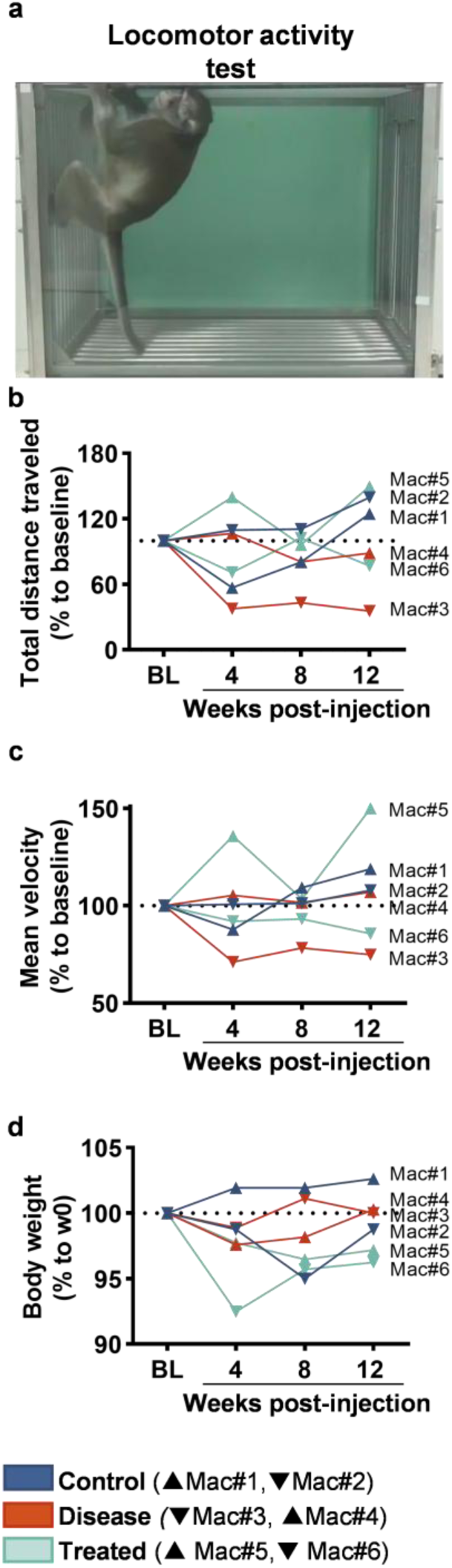
I**m**pact **of the overexpression of mutant Ataxin-3 in the cerebellum of NHP on locomotor activity. (a)** Representative image of the locomotor activity test. (**b,c**) Monkeys’ total distance travelled and mean velocity in the locomotor activity test were assessed over 30 minutes at baseline (BL), four, eight and twelve weeks after injection. Results were normalized to the BL. (**d**) Monkeys were weighed every week during the experimental period. Results are shown as a percentage of monkeys’ weight on the injection day (week zero, w0).

Animals of the control group had a total distance travelled at 12 weeks higher than at baseline. With no substantial alteration in the mean velocity, Mac#4 (disease group) exhibited a decline in the total distance travelled over the experimental period (∼11% decline at 12 weeks post-injection when compared to baseline). Likewise, the total distance travelled by Mac#3 (disease group) was significantly reduced during the whole experimental period (∼61% decline when compared to baseline). The animal with higher levels of mut*ATXN3* mRNA and Ataxin-3 aggregates (Mac#5, treated group), showed hyperactivity, with 50% more distance travelled and mean velocity at 12 weeks post-injection when compared to the baseline. On the opposite, the animal with higher levels of provirus copies, along with elevated AAV viral genomes and miRNA copies (Mac#6, treated group), showed a 23% reduction in total distance travelled and a 14% decrease in mean velocity at 12 weeks post-injection.

Throughout the experimental period, detailed weekly physical examinations were conducted to monitor the monkeys’ welfare. Hair plucking was observed in Mac#5 (treated group) before injections and continued for four weeks post-injection, while Mac#6 (treated group) ate less, moved slowly, and lost balance during the first two weeks post-injection. However, both monkeys recovered afterwards, and no additional physical alterations were observed in any other animal. Overall, body weight variations ranged from-3.85% to 2.6% relative to baseline at 12 weeks post-injection (Figure 4.d).

Due to the variability of phenotypes observed among animals overexpressing similar levels of mut*ATXN3* and the short experimental time, further studies are required to assess the effect of mutant Ataxin-3 in NHP locomotor activity.

### Overexpression of mutant Ataxin-3 in the cerebellum of NHP leads to cerebellar morphological and biochemical alterations

Despite the absence of clear motor phenotypic alterations in NHP overexpressing mut*ATXN3,* we aimed to evaluate whether any cerebellar morphological or biochemical alterations occurred.

For cerebellar morphological analysis, a brain MRI was performed at baseline, 6 and 12 weeks post-injection. Of note, due to the use of T2 MRI scans, an adjusted pipeline was created for cerebellar volumetric assessment (Figure 5.a and Supplementary Figure 3).

**Figure 5.**
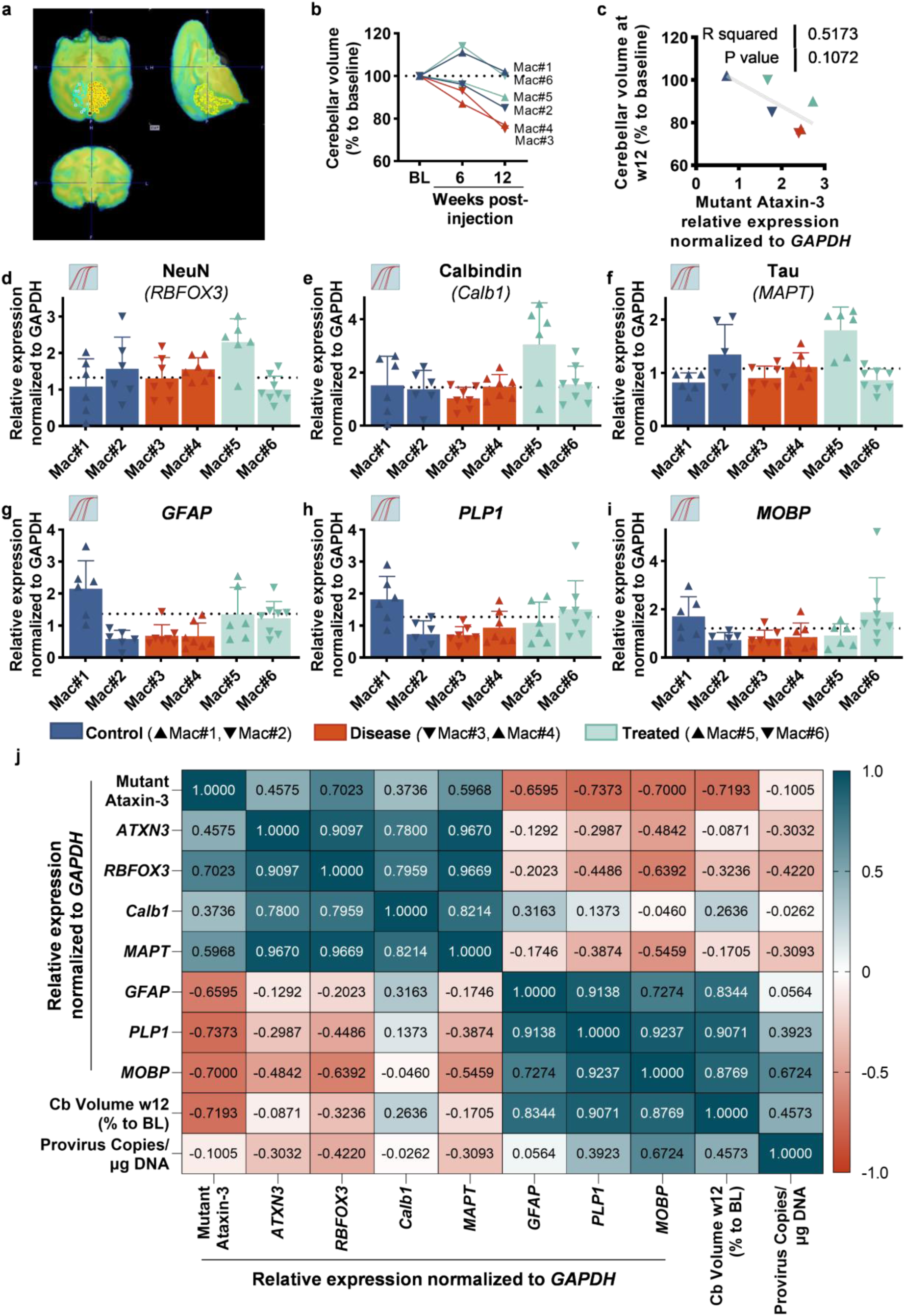
O**v**erexpression **of mutant Ataxin-3 in the cerebellum of NHP leads to cerebellar morphological and biochemical alterations. (a)** Example of the MRI analysis of cerebellar volumes. (**b**) Brain MRI was performed at BL and again at four, eight and twelve weeks post-injection. Cerebellar volume was measured at each time-point. Results are shown as a percentage of monkeys’ cerebellar volume at BL. (**c**) Simple linear regression between expression levels of mutant (human) mRNA, relative to *Macaca fascicularis GAPDH* and cerebellar volume at 12 weeks (w12) post-injection, normalised to the baseline. Data points are shown for control, disease and treated (n = 2 per group). **(d-i)** Expression levels of NeuN (*RBFOX3*) (**d**), Calbindin (Calb1) (**e**), Tau (*MAPT*) (**f**), *GAPDH* (**g**), *PLP1* (**h**) and *MOBP* (**i**) mRNA, relative to *Macaca fascicularis GAPDH*. Statistical analysis was performed using one-way ANOVA with Tukey’s post-hoc test (* *P* < 0.05, ** *P* < 0.01, *** *P* < 0.001). Results are shown as mean ± SD for control, disease, and treated (n = 5-8, per animal). The dotted line represents the average of the control group. (**j**) Multivariable correlation matrix between relative expression of different genes normalised to *Macaca fascicularis GAPDH*, provirus copies per µg of DNA and monkey’s cerebellar volume at 12 weeks (w12) post-injection, normalised to the baseline. Dark blue indicates a strong positive and dark red indicates a strong negative. The correlation was performed with control, disease and treated groups (n = 2 per group).

Although Mac#1 (control) and Mac#6 (treated) exhibited an 11-14% increase in cerebellar volume at 6 weeks post-injection, this volume returned to baseline levels by 12 weeks post-injection (Figure 5.b). In contrast, Mac#2 (control) and Mac#5 (treated) presented a decrease of 15% and 10% of the cerebellar volume at 12 weeks post-injection, respectively. Interestingly, both animals from the disease group, Mac#3 and Mac#4, showed a more pronounced decrease, with a reduction of 23% and 25% in cerebellar volume compared to the baseline, respectively.

To evaluate the relationship between mutant Ataxin-3 mRNA and cerebellar volume, a simple linear regression analysis was performed using mutant Ataxin-3 mRNA relative expression (normalised to *GAPDH*) and cerebellar volume at 12 weeks post-injection (normalised to baseline). The model showed a tendency to explain 51.73% of the variance (r^2^ = 0.5173, *p* = 0.1072) (Figure 5.c).

Next, we evaluated the mRNA levels of genes relevant to SCA3 neuropathology, including the neuronal genes *RBFOX3* (Figure 5.d), *Calb1* (Figure 5.e) and *MAPT* (Figure 5.f) and the glial genes *GFAP* (Figure 5.g), *PLP1* (Figure 5.h), and *MOBP* (Figure 5.i). Regarding neuronal genes, an increase in NeuN (*RBFOX3),* Calbindin *(Calb1)* and Tau (*MAPT*) mRNA expression was observed in the cerebellum of the treated monkey, that received the lowest dose of AAVs and that had high levels of mut*ATXN3* mRNA and Ataxin-3 aggregates (Mac#5, treated group). For glial genes (*GFAP*, *PLP1* and *MOBP*), high variability in mRNA expression was noted among animals from the control group. Nonetheless, the monkey with the highest silencing of mut*ATXN3 mRNA* (Mac#6, treated group), exhibited high levels of *PLP1* and *MOBP* mRNA when compared to the other animals expressing mut*ATXN3*.

To further understand whether the levels of mutant Ataxin-3 were associated with the observed morphological and biochemical alterations, we performed a multivariable correlation analysis (Figure 5.j and Supplementary Table 5).

Although the reduced sample size (n = 6) limited the statistical significance of the Pearson correlations (Supplementary Table 5), a strong negative correlation trend was observed between mutant Ataxin-3 mRNA levels and oligodendrocyte-related glial genes *PLP1* (r = 0.7373, *p* = 0.0945), *MOBP* (r = 0.7000, *p* = 0.1215), and a moderate negative correlation with the astrocyte-related glial gene GFAP (r = 0.6595, *p* = 0.1542). Additionally, a strong negative correlation trend was observed between mutant Ataxin-3 mRNA levels and cerebellar volume at 12 weeks post-injection (r = 0.7193, *p* = 0.1072). On the other hand, a strong positive correlation trend was noted between mutant Ataxin-3 mRNA levels and the neuronal gene *RBFOX3* (r = 0.7023, *p* = 0.1198).

Overall, mutant Ataxin-3 overexpressed in the cerebellum of NHP leads to morphological and biochemical alterations in the cerebellum that may be ameliorated by AAV9-miR-ATXN3-mediated silencing.

## Discussion

In this work, we demonstrate the following key findings: 1) Lentiviral delivery of mutant *ATXN3* to naïve NHP cerebellum induces Ataxin-3 aggregation; 2) ICM administration of AAV9-miR-ATXN3 to naïve NHP drives transgene expression in critical regions affected by SCA3 pathology; and 3) AAV9-miR-ATXN3 specifically silences mut*ATXN3* in a dose-dependent manner in the developed NHP model of SCA3.

Up to now, evidence suggests that SCA3 may have arisen from only two distinct *de novo* expansion events in the *ATXN3* gene (Kawaguchi et al., 1994). The first is thought to have occurred in Asia (TTACAC background, Joseph lineage) and the second in Portugal (GTGGCA background, Machado lineage) (Martins et al., 2007). Based on the analysis of three single nucleotide substitutions identified in linkage disequilibrium with the Ataxin-3 mutation, the exonic SNP <u>A</u>^669^TG/<u>G</u>^669^TG (rs1048755, exon 8) and <u>C</u>^987^GG/<u>G</u>^987^GG (rs12895357, exon 10) and the intronic SNP TA<u>A</u>^1118^/TA<u>C</u>^1118^ (rs3092822), four haplotypes can be found in SCA3 patients: ACA, GGC, AGA, and GGA (Gaspar et al., 2001; Goto et al., 1997; Igarashi et al., 1996; Maciel et al., 1999). Importantly, the haplotype A^669^-C^987^-A^1118^ is presented in around 72% of SCA patients and only in 2% of controls (Gaspar et al., 2001).

The affected protein, Ataxin-3, is a deubiquitinating enzyme, which is also involved in protein homeostasis, DNA damage response and chromatin organisation (reviewed in (Hernández-Carralero et al., 2024)). Some reports suggest that silencing or full abrogation of ataxin-3 does not result in significant deleterious effects (Alves et al., 2010; Boy et al., 2009; Schmitt et al., 2007; Switonski et al., 2011), but its function is still not fully understood. Indeed, Ataxin-3 is present in different tissues and cells (do Carmo Costa et al., 2004; Ichikawa et al., 2001; H. L. Paulson et al., 1997), and different conserved regions can be found across different species, suggesting that it may still have important, non-dispensable functions (Albrecht et al., 2003; Scheel, 2003). Thus, from a therapeutic standpoint, strategies that selectively silence the mutant gene and preserve the wild-type protein are preferred.

Several allele-specific gene silencing strategies for SCA3 have been developed targeting the SNP <u>C</u>^987^GG/<u>G</u>^987^GG (rs12895357), located in exon10 of the *ATXN3* gene, at the 3’end of the CAG repeat, and in linkage disequilibrium with the Ataxin-3 mutation (Alves, Hassig, et al., 2008; Conceição et al., 2016; Hauser et al., 2021; Li et al., 2004; Miller et al., 2003; Nóbrega et al., 2013, 2014; Nóbrega, Codêsso, et al., 2019). Our group recently developed an allele-specific artificial miR, delivered by an AAV9, that specifically targets this SNP (Nobre & Pereira De Almeida, 2020). This approach has been successful in achieving allele-specific silencing of mutant Ataxin-3 and recovering SCA3 pathology when administered by intravenous injection in SCA3 transgenic mice. More recently, a new construct containing two miR-*ATXN3* in tandem has been developed, to improve *ATXN3* silencing. This new bicistronic construct maintained allele specificity and achieved robust silencing of the mutant *ATXN3* allele both in vitro and in vivo. Moreover, intraparenchymal delivery and single ICM administration yielded highly promising results in different SCA3 animal models, highlighting its potential for future clinical translation (unpublished data).

However, although a wide range of SCA3 models are available, including cell models (Onofre et al., 2016), worms (Khan et al., 2006), fly (Warrick et al., 1998), fish (Watchon et al., 2017), mouse (Cemal et al., 2002) and rat (Alves, Régulier, et al., 2008) (reviewed in (Costa & Paulson, 2012; Schmidt & Schmidt, 2018)), these models have significant limitations when it comes to translating therapeutic efficacy into clinical practice, which has contributed to setbacks and failures in moving promising therapies to the clinic. With the advent of high-precision gene-targeted therapies, the development of improved animal models that more closely mimic human disorders, with similar brain structures and metabolism, has become increasingly important (Chen et al., 2023). Such models are crucial not only for evaluating the effectiveness of therapies but also for identifying disease biomarkers that can predict therapeutic outcomes and be translated into clinical practice (Tarantal et al., 2022). In 2017, Tomioka and colleagues (Tomioka, Ishibashi, et al., 2017), attempted to generate the first NHP model of SCA3. The authors delivered lentiviral vectors carrying CMV-*ATXN3*-120Q to common marmoset’ embryos. Despite considerable variability between the seven transgenic animals, those with higher expression of mutant *ATXN3* showed symptoms as early as 3 to 4 months of age. By 5.5 and 13.5 months, inclusions of Ataxin-3 and degeneration were observed in multiple regions, including the brain, but predominantly in the spinal cord, peripheral nerves, and skeletal muscles. Nevertheless, the high abortion rate (89.4%), prompted the authors to later develop a tetracycline-controllable expression system, which increased the production rate to 35.3% (Tomioka, Nogami, et al., 2017). While these SCA3 NHP models are highly valuable, generating transgenic animals in larger species is extremely challenging, as the authors documented. Additionally, maintaining transgenic colonies is very costly. The use of viral vectors for modelling neurodegenerative disorders, presents a cheaper and faster alternative, allowing for sustained transgene expression in the targeted regions (Chansel-Debordeaux & Bezard, 2019; Henriques et al., 2024).

In this work, we attempted to evaluate the feasibility of using LVs for the delivery of human mutant Ataxin-3 to NHP brains, a strategy that we have largely used in rodents for SCA3 (Alves et al., 2010; Alves, Régulier, et al., 2008; Carmona et al., 2017; Conceição et al., 2016; Cunha-Santos et al., 2016; De Almeida et al., 2002; Gonçalves et al., 2013; Mendonça et al., 2019; Nobre et al., 2022; Nóbrega et al., 2013; Rufino-Ramos et al., 2023; Santana et al., 2020; Simões et al., 2012, 2014, 2022) and has also been used in NHP for modelling Huntington’s disease (Palfi et al., 2007) and Parkison’s disease (Yang et al., 2015).

Of note, due to its wide spreading, AAVs are more commonly used for NHP disease modelling (Beckman et al., 2021; Kirik et al., 2003; Weiss et al., 2022; Yin et al., 2019); however, since we aimed to evaluate an AAV-based therapy, we opted to use LV vectors to distinguish modelling and treating within the same animal more effectively. For that, three bilateral intracerebellar LV injections were performed to maximise the area of viral transduction. Thirteen weeks after LV administration, mutant *ATXN3* mRNA and Ataxin-3 aggregates were detected in the NHP cerebellum. These Ataxin-3 aggregates appear to be neurotoxic, as they led to the recruitment of inflammatory cells to the Ataxin-3 aggregates region. Higher levels of mutant *ATXN3* were also associated with an increase in the neuronal markers *NeuN*, *Calb1* and *MAPT*, which we hypothesise to be a compensatory response to the cellular stress caused by the presence of Ataxin-3 aggregates (Dell’Orco et al., 2015; Kim & Suh, 2022). In a future study, it would be fundamental to evaluate the overexpression of mutant *ATXN3* in the cerebellum of NHP over longer time points, to assess if exogenous mut*ATXN3* can trigger further neurodegeneration and neuropathology, or if the primate immune system will be able to clear mutant ATXN3 and lentiviral transduced cells. Moreover, evaluating different doses of LV vectors would also be important. Notwithstanding, the successful overexpression of mut*ATXN3* in the cerebellum of *Macaca fascicularis* mediated by LV delivery offers a new powerful tool for the evaluation of novel therapies for SCA3.

Next, we aimed to evaluate a previously developed AAV-based strategy for the silencing of Ataxin-3 in this newly developed NHP SCA3 model. Due to its safety profile and sustained transgene expression, AAVs are the gene delivery vectors of choice for the central nervous system (CNS) (Saraiva et al., 2016). Among the seven AAV products approved for clinical use, there are already three products for the treatment of nervous system disorders: Onasemnogene abeparvovec (Zolgensma) for Spinal muscular atrophy (SMA), approved in 2019, in which AAV9-h*SMN* (human survival motor neuron gene) is delivered IV (Mahajan, 2019; Mendell et al., 2021); Eladocagene exuparvovec (Upstaza) for Aromatic L-amino acid decarboxylase (AADC) deficiency, approved in 2022, in which four intraputaminal administrations of AAV2-hDDC (human dopa decarboxylase) are performed (Keam, 2022; Tai et al., 2022); and Delandistrogene moxeparvovec-rokl (Elevidys) for Duchenne muscular dystrophy (DMD), approved in 2023, in which AAVrh74-micro-dystrophin is administered IV (Hoy, 2023; Mendell et al., 2020). Due to its minimal invasiveness, IV administration is a highly attractive therapeutic route; however, high doses of vectors are required, which can result in off-target effects in peripheral organs, such as the liver and the heart and neutralisation by anti-AAV9 antibodies (Huang et al., 2021; Louis Jeune et al., 2013; Prasad et al., 2022). To overcome these limitations, an intra-CSF approach, in which AAVs are directly infused into the CSF, reducing peripheral transduction while circumventing the BBB, has recently gained a lot of interest in preclinical and clinical studies (Chen et al., 2023; Marchi et al., 2022). Intra-CSF delivery of AAVs can be achieved through intracerebroventricular (ICV), ICM and intra-thecal (IT) administrations (Perez et al., 2020). Each of these approaches results in distinct patterns of CNS transduction, with ICM administration of AAV9 being able to achieve a widespread transgene delivery throughout the CNS, with minimal transduction of peripheral organs (Marchi et al., 2022).

In the present study, to evaluate the biodistribution of our silencing strategy, a low (0.3 x 10^13^ vg, d1) and high (3.0 x 10^13^ vg, d2) dose of AAV encoding an allele-specific artificial miR against mutant *ATXN3* mRNA were ICM-delivered to naïve *Macaca fascicularis*. As previously described in other studies, viral genomes could be found in the whole brain, with the highest transduction observed in the SC and DRG (Gray et al., 2013; Hinderer et al., 2014; Hordeaux et al., 2019; Samaranch et al., 2013; Wiseman et al., 2024). Accordingly, artificial miRNA copies were primarily detected in the hindbrain, SC, and DRG. No adverse events were observed and no differences in the levels of the astrocytic marker GFAP were detected in any of the animals. Since AAV9-miR-ATXN3 delivered by ICM injection reached the most degenerated brain regions in SCA3 without causing side effects or astrogliosis at either dose, we proceeded to test AAV-miR-ATXN3 target engagement in SCA3 NHPs using a dose of 2.0 x 10^13^ vg per animal (2.5 x 10^12^ ± 0.33 x 10^12^ vg per kg of body weight).

In the AAV-miR-ATXN3 target engagement experiment, treated Mac#5 exhibited 13.6-fold fewer proviral copies per microgram of DNA compared to treated Mac#6, and its AAV viral genomes were close to the detection limit. As expected, no reduction in mutant *ATXN3* mRNA or aggregates was observed in this animal compared to those in the disease group. On the other hand, treated Mac#6, which had higher levels of provirus copies, but also higher levels of AAV viral genomes and artificial miRNA copies, showed a reduction of mutant *ATXN3* mRNA, and no ATXN3 aggregates were detected in the cerebellum. Moreover, treated Mac#6 had higher levels of myelin-related proteins, *PLP1* and *MOBP*, compared to animals with higher levels of mutant *ATXN3*. These findings suggest that the presence of mutant *ATXN3* is accompanied by a reduction of these glial proteins, as has been observed in other SCA3 animal models (Schuster et al., 2022). Importantly, these alterations appeared to be reverted by the AAV9-miR-ATXN3 treatment. No alterations were observed in endogenous *ATXN3* mRNA and protein. Despite further studies being required, AAV9-miR-ATXN3 seems to be able to specifically silence mut*ATXN3* in a dose-dependent manner in our newly developed NHP SCA3 model.

This pilot study was underpowered to draw definitive conclusions as there was great variability among the animals in each study group. Table 1 provides a summary of all the main observations for each animal. Unlike commonly used animal models, NHP are outbred, meaning they have different genetic backgrounds, similar to humans. As a result, they exhibit greater physiological and phenotypic variability, which can deceive the outcomes but make them more predictive of the clinical scenario (Vallender & Miller, 2013). Notwithstanding, the findings from this work are crucial, not only for advancing AAV9-miRNA-ATXN3 toward clinical use but also for establishing a valuable platform for validating future biomarkers or therapeutic strategies.

**Table 1.**
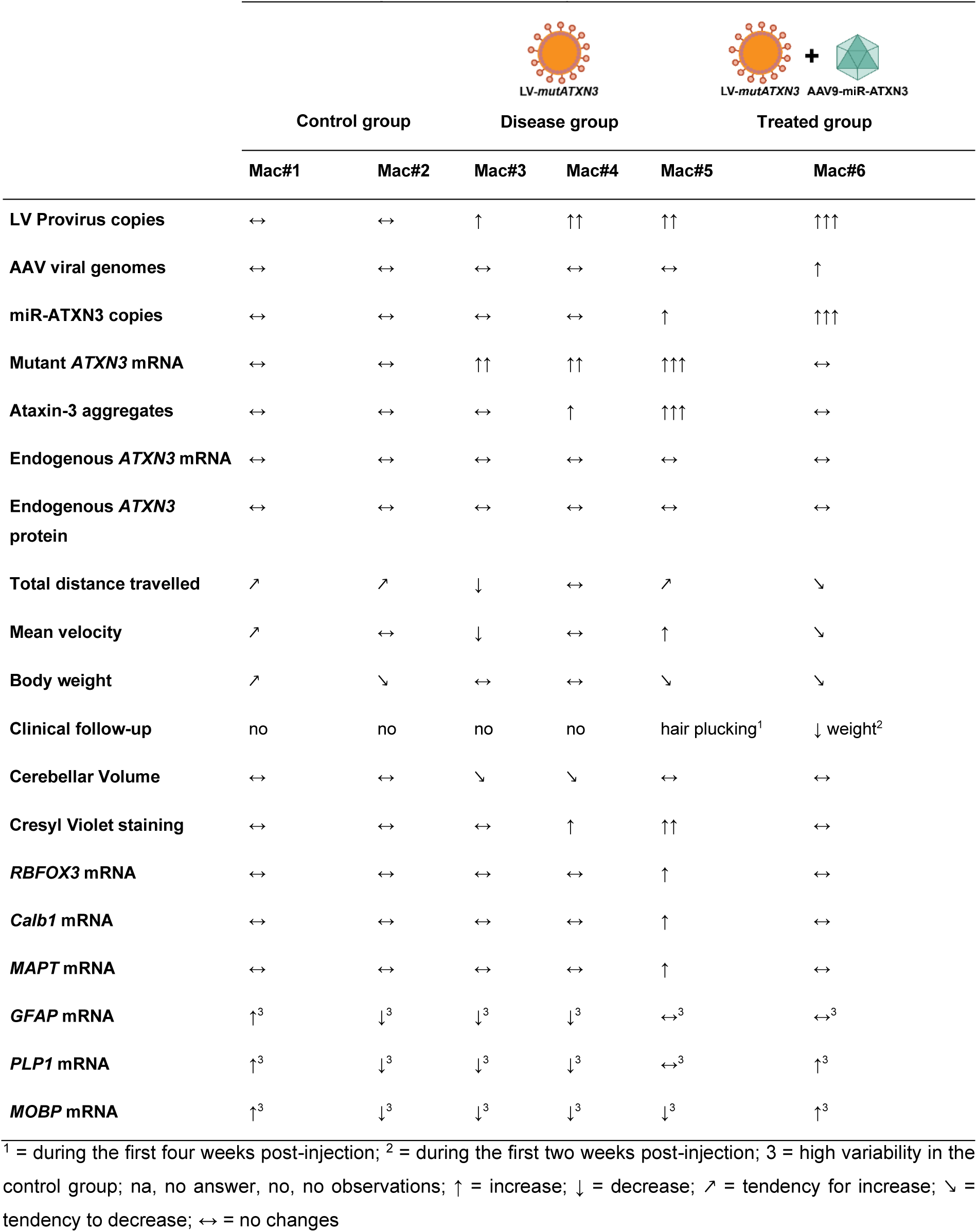
Overview of th e main findings for each monkey.

|  |  |  | LV-mutATXN3 |  | LV-mutATXN3 + AAV9-miR-ATXN3 |  |
| --- | --- | --- | --- | --- | --- | --- |
|  | Control group |  | Disease group |  | Treated group |  |
|  | Mac#1 | Mac#2 | Mac#3 | Mac#4 | Mac#5 | Mac#6 |
| LV Provirus copies | ↔ | ↔ | ↑ | ↑↑ | ↑↑ | ↑↑↑ |
| AAV viral genomes | ↔ | ↔ | ↔ | ↔ | ↔ | ↑ |
| miR-ATXN3 copies | ↔ | ↔ | ↔ | ↔ | ↑ | ↑↑↑ |
| Mutant *ATXN3* mRNA | ↔ | ↔ | ↑↑ | ↑↑ | ↑↑↑ | ↔ |
| Ataxin-3 aggregates | ↔ | ↔ | ↔ | ↑ | ↑↑↑ | ↔ |
| Endogenous *ATXN3* mRNA | ↔ | ↔ | ↔ | ↔ | ↔ | ↔ |
| Endogenous *ATXN3* protein | ↔ | ↔ | ↔ | ↔ | ↔ | ↔ |
| Total distance travelled | ↗ | ↗ | ↓ | ↔ | ↗ | ↘ |
| Mean velocity | ↗ | ↔ | ↓ | ↔ | ↑ | ↘ |
| Body weight | ↗ | ↘ | ↔ | ↔ | ↘ | ↘ |
| Clinical follow-up | no | no | no | no | hair plucking<sup>1</sup> | ↓ weight<sup>2</sup> |
| Cerebellar Volume | ↔ | ↔ | ↘ | ↘ | ↔ | ↔ |
| Cresyl Violet staining | ↔ | ↔ | ↔ | ↑ | ↑↑ | ↔ |
| *RBFOX3* mRNA | ↔ | ↔ | ↔ | ↔ | ↑ | ↔ |
| *Calb1* mRNA | ↔ | ↔ | ↔ | ↔ | ↑ | ↔ |
| *MAPT* mRNA | ↔ | ↔ | ↔ | ↔ | ↑ | ↔ |
| *GFAP* mRNA | ↑<sup>3</sup> | ↓<sup>3</sup> | ↓<sup>3</sup> | ↓<sup>3</sup> | ↔<sup>3</sup> | ↔<sup>3</sup> |
| *PLP1* mRNA | ↑<sup>3</sup> | ↓<sup>3</sup> | ↓<sup>3</sup> | ↓<sup>3</sup> | ↔<sup>3</sup> | ↑<sup>3</sup> |
| *MOBP* mRNA | ↑<sup>3</sup> | ↓<sup>3</sup> | ↓<sup>3</sup> | ↓<sup>3</sup> | ↓<sup>3</sup> | ↑<sup>3</sup> |

<sup>1</sup> = during the first four weeks post-injection; <sup>2</sup> = during the first two weeks post-injection; 3 = high variability in the control group; na, no answer, no, no observations; ↑ = increase; ↓ = decrease; ↗ = tendency for increase; ↘ = tendency to decrease; ↔ = no changes

## Author Contributions

C.H., P.Y, S.d.M, R.A.B., L.P.d.A. and R.J.N. conceived and designed the experiments. C.H., A.F., C.J., M.G., J.F. and R.A.B. performed the experiments. C.H. analysed the data. J.C., assisted with MRI data analysis. D.L., A.C.S., A.R.F., M.M.L., J.C., M.C.B., P.Y, S.d.M, and P.H, shared knowledge. C.H. drafted the initial version of the manuscript. All authors reviewed and contributed to editing the final version.

## Funding and ackowledgments

This work was partially funded by PTC Therapeutics. Our group is also supported by the European Regional Development Fund (ERDF) through the Centro 2020 Regional Operational Programme, the Operational Programme for Competitiveness and Internationalisation (COMPETE 2020), and by Portuguese national funds via Fundação para a Ciência e a Tecnologia (FCT), under the projects UIDB/04539/2025, UIDP/04539/2025, LA/P/0058/2020 and 2022.06118.PTDC; as well as by the projects SpreadSilencing (POCI-01-0145-FEDER029716), ViraVector (CENTRO-01-0145-FEDER-022095) and Neurodiet (JPND/0001/2022). Additional support was provided by CinTech under PRR, ARDAT under the IMI2 JU Grant agreement No 945473 (co-funded by EU and EFPIA), and Capacity 2023 (ID: 101145599), GeneT (ID: 101059981), ERDERA (ID: 101156595), GeneH (ID: 101186939) and GCure (ID: 101186929), under the European Union’s Horizon Europe program. Further funding was received from the American Portuguese Biomedical Research Fund (APBRF), the European Advanced Translational Research Infrastructure for Neurosciences (NeurATRIS), and the Richard Chin and Lily Lock MJD Research Fund. FCT PhD fellowships supported the work of C.H. (2021.06939.BD), D.L. (2020.09668.BD), A.C.S. (2020.07721.BD), A.R.F. (2021.08215.BD), and M.M.L. (2021.05776.BD). We also thank all members of the L.P.A. laboratory for the support and discussions.

## Competing interests

R.J.N. and L.P.A. are inventors on patent application WO2020144611A1, which is related to the subject of this manuscript. P.Y. is employee of PTC Therapeutics. S.d.M. was an employee of PTC Therapeutics during the course of this study. All other authors declare no competing interests.

**See supplemental material for supplementary figures and tables**

## References

Albrecht, M., Hoffmann, D., Evert, B. O., Schmitt, I., Wüllner, U., & Lengauer, T. (2003). Structural modeling of ataxin-3 reveals distant homology to adaptins. Proteins: Structure, Function, and Bioinformatics, 50(2), 355–370. 10.1002/prot.10280

Alves, S., Hassig, R., Brouillet, E., Lima, M. C. P. D., Hantraye, P., Nascimento-Ferreira, I., Auregan, G., Hassig, R., Dufour, N., Brouillet, E., Pedroso de Lima, M. C., Hantraye, P., de Almeida, L. P., & Déglon, N. (2008). Allele-specific RNA silencing of mutant ataxin-3 mediates neuroprotection in a rat model of Machado-Joseph disease. PLoS ONE, 3(10). 10.1371/journal.pone.0003341

Alves, S., Nascimento-Ferreira, I., Dufour, N., Hassig, R., Auregan, G., Nóbrega, C., Brouillet, E., Hantraye, P., Pedroso de Lima, M. C., Déglon, N., & de Almeida, L. P. (2010). Silencing ataxin-3 mitigates degeneration in a rat model of Machado-Joseph disease: No role for wild-type ataxin-3? Human Molecular Genetics, 19(12), 2380–2394. 10.1093/hmg/ddq111

Alves, S., Régulier, E., Nascimento-Ferreira, I., Hassig, R., Dufour, N., Koeppen, A., Carvalho, A. L., Simões, S., De Lima, M. C. P., Brouillet, E., Gould, V. C., Déglon, N., & De Almeida, L. P. (2008). Striatal and nigral pathology in a lentiviral rat model of Machado-Joseph disease. Human Molecular Genetics, 17(14), 2071–2083. 10.1093/hmg/ddn106

Beckman, D., Chakrabarty, P., Ott, S., Dao, A., Zhou, E., Janssen, W. G., Donis-Cox, K., Muller, S., Kordower, J. H., & Morrison, J. H. (2021). A novel tau-based rhesus monkey model of Alzheimer’s pathogenesis. Alzheimer’s and Dementia, 17(6), 933–945. 10.1002/alz.12318

Bettencourt, C., & Lima, M. (2011). Machado-Joseph disease: From first descriptions to new perspectives. Orphanet Journal of Rare Diseases, 6(1), 35. 10.1186/1750-1172-6-35

Boy, J., Schmidt, T., Wolburg, H., Mack, A., Nuber, S., Böttcher, M., Schmitt, I., Holzmann, C., Zimmermann, F., Servadio, A., & Riess, O. (2009). Reversibility of symptoms in a conditional mouse model of spinocerebellar ataxia type 3. Human Molecular Genetics, 18(22), 4282–4295. 10.1093/hmg/ddp381

Carlos, J., Belmonte, I., Callaway, E. M., Churchland, P., Sarah, J., Feng, G., Homanics, G. E., Lee, K.-F., Leopold, D. A., Cory, T., Mitchell, J. F., Mitalipov, S., Moutri, A. R., Anthony, J., Izpisua Belmonte, J. C., Callaway, E. M., Caddick, S. J., Churchland, P., Feng, G.,… Zhang, F. (2016). Brains, Genes and Primates Juan. Neuron, 86(3), 617– 631. 10.1016/j.neuron.2015.03.021.Brains

Carmona, V., Cunha-Santos, J., Onofre, I., Simões, A. T., Vijayakumar, U., Davidson, B. L., & Pereira de Almeida, L. (2017). Unravelling Endogenous MicroRNA System Dysfunction as a New Pathophysiological Mechanism in Machado-Joseph Disease. Molecular Therapy, 25(4), 1038–1055. 10.1016/j.ymthe.2017.01.021

Cemal, C. K., Carroll, C. J., Lawrence, L., Lowrie, M. B., Ruddle, P., Al-Mahdawi, S., King, R. H. M., Pook, M. A., Huxley, C., & Chamberlain, S. (2002). YAC transgenic mice carrying pathological alleles of the MJD1 locus exhibit a mild and slowly progressive cerebellar deficit. Human Molecular Genetics, 11(9), 1075–1094. 10.1093/hmg/11.9.1075

Chai, Y., Koppenhafer, S. L., Bonini, N. M., & Paulson, H. L. (1999). Analysis of the Role of Heat Shock Protein (Hsp) Molecular Chaperones in Polyglutamine Disease. The Journal of Neuroscience, 19(23), 10338. 10.1523/JNEUROSCI.19-23-10338.1999

Chansel-Debordeaux, L., & Bezard, E. (2019). Local transgene expression and whole-body transgenesis to model brain diseases in nonhuman primate. Animal Models and Experimental Medicine, 2(1), 9–17. 10.1002/ame2.12055

Chen, X., Lim, D. A., Lawlor, M. W., Dimmock, D., Vite, C. H., Lester, T., Tavakkoli, F., Sadhu, C., Prasad, S., & Gray, S. J. (2023). Biodistribution of Adeno-Associated Virus Gene Therapy Following Cerebrospinal Fluid-Directed Administration. Human Gene Therapy, 34(3–4), 94–111. 10.1089/hum.2022.163

Conceição, M., Costa, P., Conceiç, M., Hirai, H., Pereira, L., Almeida, D., Mendonça, L., Nóbrega, C., Gomes, C., Costa, P., Hirai, H., Moreira, J. N., Lima, M. C., Manjunath, N., & Pereira de Almeida, L. (2016). Intravenous administration of brain-targeted stable nucleic acid lipid particles alleviates Machado-Joseph disease neurological phenotype. Biomaterials, 82, 124–137. 10.1016/j.biomaterials.2015.12.021

Costa, M. do C., & Paulson, H. L. (2012). Toward understanding Machado-Joseph Disease. Progress in Neurobiology, 97(2), 239–257. 10.1016/j.pneurobio.2011.11.006

Cunha-Santos, J., Duarte-Neves, J., Carmona, V., Guarente, L., De Almeida, L. P., & Cavadas, C. (2016). Caloric restriction blocks neuropathology and motor deficits in Machado-Joseph disease mouse models through SIRT1 pathway. Nature Communications, 7, 1–14. 10.1038/ncomms11445

Datson, N. A., González-Barriga, A., Kourkouta, E., Weij, R., van de Giessen, J., Mulders, S., Kontkanen, O., Heikkinen, T., Lehtimäki, K., & van Deutekom, J. C. T. (2017). The expanded CAG repeat in the huntingtin gene as target for therapeutic RNA modulation throughout the HD mouse brain. PloS One, 12(2), e0171127. 10.1371/journal.pone.0171127

De Almeida, L. P., Ross, C. A., Zala, D., Aebischer, P., & Déglon, N. (2002). Lentiviral-Mediated Delivery of Mutant Huntingtin in the Striatum of Rats Induces a Selective Neuropathology Modulated by Polyglutamine Repeat Size, Huntingtin Expression Levels, and Protein Length. Journal of Neuroscience, 22(9). 10.1523/jneurosci.22-09-03473.2002

de Sousa-Lourenço, J., Silva, A. C., Pereira de Almeida, L., & Nobre, R. J. (2024). Molecular therapy for polyQ disorders: From bench to clinical trials. Trends in Molecular Medicine, 30(9), 804–808. 10.1016/j.molmed.2024.05.004

Dell’Orco, J. M., Wasserman, A. H., Chopra, R., Ingram, M. A. C., Hu, Y.-S., Singh, V., Wulff, H., Opal, P., Orr, H. T., & Shakkottai, V. G. (2015). Neuronal Atrophy Early in Degenerative Ataxia Is a Compensatory Mechanism to Regulate Membrane Excitability. The Journal of Neuroscience, 35(32), 11292. 10.1523/JNEUROSCI.1357-15.2015

do Carmo Costa, M., Gomes-da-Silva, J., Miranda, C. J., Sequeiros, J., Santos, M. M., & Maciel, P. (2004). Genomic structure, promoter activity, and developmental expression of the mouse homologue of the Machado–Joseph disease (*MJD*) gene. Genomics, 84(2), 361–373. 10.1016/j.ygeno.2004.02.012

Duarte-Neves, J., Gonçalves, N., Cunha-Santos, J., Simões, A. T., den Dunnen, W. F. A., Hirai, H., Kügler, S., Cavadas, C., & Pereira de Almeida, L. (2015). Neuropeptide Y mitigates neuropathology and motor deficits in mouse models of Machado–Joseph disease. Human Molecular Genetics, 24(19), 5451–5463. 10.1093/hmg/ddv271

Evers, M. M., Pepers, B. A., Deutekom, J. C. T. van, Mulders, S. A. M., Dunnen, J. T. den, Aartsma-Rus, A., Ommen, G.-J. B. van, & Roon-Mom, W. M. C. van. (2011). Targeting Several CAG Expansion Diseases by a Single Antisense Oligonucleotide. PLOS ONE, 6(9), e24308. 10.1371/journal.pone.0024308

Fardghassemi, Y., Maios, C., & Parker, J. A. (2021). Small Molecule Rescue of ATXN3 Toxicity in C. elegans via TFEB/HLH-30. Neurotherapeutics, 18(2), 1151–1165. 10.1007/s13311-020-00993-5

Gaspar, C., Lopes-Cendes, I., Hayes, S., Goto, J., Arvidsson, K., Dias, A., Silveira, I., Maciel, P., Coutinho, P., Lima, M., Zhou, Y.-X., Soong, B.-W., Watanabe, M., Giunti, P., Stevanin, G., Riess, O., Sasaki, H., Hsieh, M., Nicholson, G. A.,… Rouleau, G. A. (2001). Ancestral Origins of the Machado-Joseph Disease Mutation: A Worldwide Haplotype Study. American Journal of Human Genetics, 68(2), 523–528.

Gonçalves, N., Simões, A. T., Cunha, R. A., & de Almeida, L. P. (2013). Caffeine and adenosine A(2A) receptor inactivation decrease striatal neuropathology in a lentiviral-based model of Machado-Joseph disease. Annals of Neurology, 73(5), 655–666. 10.1002/ana.23866

Goto, J., Watanabe, M., Ichikawa, Y., Yee, S.-B., Ihara, N., Endo, K., Igarashi, S., Takiyama, Y., Gaspar, C., Maciel, P., Tsuji, S., Rouleau, G. A., & Kanazawa, I. (1997). Machado– Joseph disease gene products carrying different carboxyl termini. Neuroscience Research, 28(4), 373–377. 10.1016/S0168-0102(97)00056-4

Gray, S. J., Nagabhushan Kalburgi, S., McCown, T. J., & Jude Samulski, R. (2013). Global CNS gene delivery and evasion of anti-AAV-neutralizing antibodies by intrathecal AAV administration in non-human primates. Gene Therapy, 20(4), 450–459. 10.1038/gt.2012.101

Haas, E., Incebacak, R. D., Hentrich, T., Huridou, C., Schmidt, T., Casadei, N., Maringer, Y., Bahl, C., Zimmermann, F., Mills, J. D., Aronica, E., Riess, O., Schulze-Hentrich, J. M., & Hübener-Schmid, J. (2022). A Novel SCA3 Knock-in Mouse Model Mimics the Human SCA3 Disease Phenotype Including Neuropathological, Behavioral, and Transcriptional Abnormalities Especially in Oligodendrocytes. Molecular Neurobiology, 59(1), 495–522. 10.1007/s12035-021-02610-8

Harding, J. D. (2013). Progress in genetics and genomics of nonhuman primates. ILAR Journal, 54(2), 77–81. 10.1093/ilar/ilt051

Hauser, S., Helm, J., Kraft, M., Korneck, M., Hübener-Schmid, J., & Schöls, L. (2021). Allele-specific targeting of mutant ataxin-3 by antisense oligonucleotides in SCA3-iPSC-derived neurons. Molecular Therapy. Nucleic Acids, 27, 99–108. 10.1016/j.omtn.2021.11.015

He, L., Wang, S., Peng, L., Zhao, H., Li, S., Han, X., Habimana, J. de D., Chen, Z., Wang, C., Peng, Y., Peng, H., Xie, Y., Lei, L., Deng, Q., Wan, L., Wan, N., Yuan, H., Gong, Y., Zou, G.,… Jiang, H. (2021). CRISPR/Cas9 mediated gene correction ameliorates abnormal phenotypes in spinocerebellar ataxia type 3 patient-derived induced pluripotent stem cells. Translational Psychiatry, 11(1), 1–13. 10.1038/s41398-021-01605-2

Henriques, C., Lopes, M. M., Silva, A. C., Lobo, D. D., Badin, R. A., Hantraye, P., Pereira de Almeida, L., & Nobre, R. J. (2024). Viral-based animal models in polyglutamine disorders. Brain, 147(4), 1166–1189. 10.1093/brain/awae012

Hernández-Carralero, E., Quinet, G., & Freire, R. (2024). ATXN3: A multifunctional protein involved in the polyglutamine disease spinocerebellar ataxia type 3. Expert Reviews in Molecular Medicine, 26, e19. 10.1017/erm.2024.10

Hinderer, C., Bell, P., Vite, C. H., Louboutin, J. P., Grant, R., Bote, E., Yu, H., Pukenas, B., Hurst, R., & Wilson, J. M. (2014). Widespread gene transfer in the central nervous system of cynomolgus macaques following delivery of AAV9 into the cisterna magna. Molecular Therapy - Methods and Clinical Development, 1(August), 14051. 10.1038/mtm.2014.51

Hordeaux, J., Hinderer, C., Buza, E. L., Louboutin, J.-P., Jahan, T., Bell, P., Chichester, J. A., Tarantal, A. F., & Wilson, J. M. (2019). Safe and Sustained Expression of Human Iduronidase After Intrathecal Administration of Adeno-Associated Virus Serotype 9 in Infant Rhesus Monkeys. Human Gene Therapy, 30(8), 957–966. 10.1089/hum.2019.012

Hoy, S. M. (2023). Delandistrogene Moxeparvovec: First Approval. Drugs, 83(14), 1323–1329. 10.1007/s40265-023-01929-x

Hu, J., Gagnon, K. T., Liu, J., Watts, J. K., Syeda-Nawaz, J., Bennett, C. F., Swayze, E. E., Randolph, J., Chattopadhyaya, J., & Corey, D. R. (2011). Allele-selective inhibition of ataxin-3 (ATX3) expression by antisense oligomers and duplex RNAs. 392(4), 315–325. 10.1515/bc.2011.045

Hu, J., Matsui, M., Gagnon, K. T., Schwartz, J. C., Gabillet, S., Arar, K., Wu, J., Bezprozvanny, I., & Corey, D. R. (2009). Allele-specific silencing of mutant huntingtin and ataxin-3 genes by targeting expanded CAG repeats in mRNAs. Nature Biotechnology, 27(5), 478–484. 10.1038/nbt.1539

Huang, L., Wan, J., Wu, Y., Tian, Y., Yao, Y., Yao, S., Ji, X., Wang, S., Su, Z., & Xu, H. (2021). Challenges in adeno-associated virus-based treatment of central nervous system diseases through systemic injection. Life Sciences, 270. 10.1016/J.LFS.2021.119142

Ichikawa, Y., Goto, J., Hattori, M., Toyoda, A., Ishii, K., Jeong, S.-Y., Hashida, H., Masuda, N., Ogata, K., Kasai, F., Hirai, M., Maciel, P., Rouleau, G. A., Sakaki, Y., & Kanazawa, I. (2001). The genomic structure and expression of MJD, the Machado-Joseph disease gene. Journal of Human Genetics, 46(7), 413–422. 10.1007/s100380170060

Igarashi, S., Takiyama, Y., Cancel, G., Rogaeva, E. A., Sasaki, H., Wakisaka, A., Zhou, Y. X., Takano, H., Endo, K., Sanpei, K., Oyake, M., Tanaka, H., Stevanin, G., Abbas, N., Dürr, A., Rogaev, E. I., Sherrington, R., Tsuda, T., Ikeda, M.,… Tsuji, S. (1996). Intergenerational instability of the CAG repeat of the gene for Machado-Joseph disease (MJD1) is affected by the genotype of the normal chromosome: Implications for the molecular mechanisms of the instability of the CAG repeat. Human Molecular Genetics, 5(7), 923–932. 10.1093/hmg/5.7.923

Kawaguchi, Y., Okamoto, T., Taniwaki, M., Aizawa, M., Inoue, M., Katayama, S., Kawakami, H., Nakamura, S., Nishimura, M., Akiguchi, I., Kimura, J., Narumiya, S., & Kakizuka, A. (1994). CAG expansions in a novel gene for Machado-Joseph disease at chromosome 14q32.1. Nature Genetics, 8(3), 221–228. 10.1038/ng1194-221

Keam, S. J. (2022). Eladocagene Exuparvovec: First Approval. Drugs, 82(13), 1427–1432. 10.1007/s40265-022-01775-3

Khan, L. A., Bauer, P. O., Miyazaki, H., Lindenberg, K. S., Landwehrmeyer, B. G., & Nukina, N. (2006). Expanded polyglutamines impair synaptic transmission and ubiquitin– proteasome system in Caenorhabditis elegans. Journal of Neurochemistry, 98(2), 576–587. 10.1111/j.1471-4159.2006.03895.x

Kieling, C., Prestes, P. R., Saraiva-Pereira, M. L., & Jardim, L. B. (2007). Survival estimates for patients with Machado-Joseph disease (SCA3). Clinical Genetics, 72(6), 543–545. 10.1111/j.1399-0004.2007.00910.x

Kim, Y., & Suh, B.-C. (2022). Editorial: Brain cells’ compensatory mechanisms in response to disease risk factors. Frontiers in Molecular Neuroscience, 15. 10.3389/fnmol.2022.1096287

Kirik, D., Annett, L. E., Burger, C., Muzyczka, N., Mandel, R. J., & Björklund, A. (2003). Nigrostriatal α-synucleinopathy induced by viral vector-mediated overexpression of human α-synuclein: A new primate model of Parkinson’s disease. Proceedings of the National Academy of Sciences of the United States of America, 100(5), 2884–2889. 10.1073/pnas.0536383100

Klockgether, T., Mariotti, C., & Paulson, H. L. (2019). Spinocerebellar ataxia. Nature Reviews Disease Primers, 5(1), 1–21. 10.1038/s41572-019-0074-3

Koppenol, R., Conceição, A., Afonso, I. T., Afonso-Reis, R., Costa, R. G., Tomé, S., Teixeira, D., da Silva, J. P., Côdesso, J. M., Brito, D. V. C., Mendonça, L., Marcelo, A., Pereira de Almeida, L., Matos, C. A., & Nóbrega, C. (2023). The stress granule protein G3BP1 alleviates spinocerebellar ataxia-associated deficits. Brain: A Journal of Neurology, 146(6), 2346–2363. 10.1093/brain/awac473

Kourkouta, E., Weij, R., González-Barriga, A., Mulder, M., Verheul, R., Bosgra, S., Groenendaal, B., Puoliväli, J., Toivanen, J., van Deutekom, J. C. T., & Datson, N. A. (2019). Suppression of Mutant Protein Expression in SCA3 and SCA1 Mice Using a CAG Repeat-Targeting Antisense Oligonucleotide. Molecular Therapy. Nucleic Acids, 17, 601–614. 10.1016/j.omtn.2019.07.004

Kumar, S., & Hedges, S. B. (1998). A molecular timescale for vertebrate evolution. Nature, 392(6679), 917–920. 10.1038/31927

La Spada, A. R., & Taylor, J. P. (2003). Polyglutamines placed into context. Neuron, 38(5), 681–684. 10.1016/S0896-6273(03)00328-3

Lee, B.-H., Lee, M. J., Park, S., Oh, D.-C., Elsasser, S., Chen, P.-C., Gartner, C., Dimova, N., Hanna, J., Gygi, S. P., Wilson, S. M., King, R. W., & Finley, D. (2010). Enhancement of proteasome activity by a small-molecule inhibitor of USP14. Nature, 467(7312), 179–184. 10.1038/nature09299

Li, Y., Yokota, T., Matsumura, R., Taira, K., & Mizusawa, H. (2004). Sequence-dependent and independent inhibition specific for mutant ataxin-3 by small interfering RNA. Annals of Neurology, 56(1), 124–129. 10.1002/ana.20141

Lin, T.-H., Chen, W.-L., Hsu, S.-F., Chen, I.-C., Lin, C.-H., Chang, K.-H., Wu, Y.-R., Chen, Y.-R., Yao, C.-F., Lin, W., Lee-Chen, G.-J., & Chen, C.-M. (2024). Small Molecules Inducing Autophagic Degradation of Expanded Polyglutamine Protein through Interaction with Both Mutant ATXN3 and LC3. International Journal of Molecular Sciences, 25(19), Article 19. 10.3390/ijms251910707

Louis Jeune, V., Joergensen, J. A., Hajjar, R. J., & Weber, T. (2013). Pre-existing Anti–Adeno-Associated Virus Antibodies as a Challenge in AAV Gene Therapy. Human Gene Therapy Methods, 24(2), 59–67. 10.1089/hgtb.2012.243

Maciel, P., Gaspar, C., Guimarães, L., Goto, J., Lopes-Cendes, I., Hayes, S., Arvidsson, K., Dias, A., Sequeiros, J., Sousa, A., & Rouleau, G. A. (1999). Study of three intragenic polymorphisms in the Machado-Joseph disease gene (MJD1) in relation to genetic instability of the (CAG)n tract. European Journal of Human Genetics, 7(2), 147–156. 10.1038/sj.ejhg.5200264

Mahajan, R. (2019). Onasemnogene Abeparvovec for Spinal Muscular Atrophy: The Costlier Drug Ever. International Journal of Applied and Basic Medical Research, 9(3), 127. 10.4103/ijabmr.IJABMR_190_19

Marchi, P. M., Marrone, L., & Azzouz, M. (2022). Delivery of therapeutic AAV9 vectors via cisterna magna to treat neurological disorders. Trends in Molecular Medicine, 28(1), 79–80. 10.1016/j.molmed.2021.09.007

Martier, R., Sogorb-Gonzalez, M., Stricker-Shaver, J., Hübener-Schmid, J., Keskin, S., Klima, J., Toonen, L. J., Juhas, S., Juhasova, J., Ellederova, Z., Motlik, J., Haas, E., van Deventer, S., Konstantinova, P., Nguyen, H. P., & Evers, M. M. (2019). Development of an AAV-Based MicroRNA Gene Therapy to Treat Machado-Joseph Disease. Molecular Therapy - Methods and Clinical Development, 15(December), 343–358. 10.1016/j.omtm.2019.10.008

Martins, S., Calafell, F., Gaspar, C., Wong, V. C. N., Silveira, I., Nicholson, G. A., Brunt, E. R., Tranebjaerg, L., Stevanin, G., Hsieh, M., Soong, B.-W., Loureiro, L., Dürr, A., Tsuji, S., Watanabe, M., Jardim, L. B., Giunti, P., Riess, O., Ranum, L. P. W.,… Sequeiros, J. (2007). Asian origin for the worldwide-spread mutational event in Machado-Joseph disease. Archives of Neurology, 64(10), 1502–1508. 10.1001/archneur.64.10.1502

Matos, C. A., Carmona, V., Vijayakumar, U. G., Lopes, S., Albuquerque, P., Conceição, M., Nobre, R. J., Nóbrega, C., & de Almeida, L. P. (2018). Gene therapies for polyglutamine diseases. In Advances in Experimental Medicine and Biology (Vol. 1049, pp. 395–438). 10.1007/978-3-319-71779-1_20

McLoughlin, H. S., Gundry, K., Rainwater, O., Schuster, K. H., Wellik, I. G., Zalon, A. J., Benneyworth, M. A., Eberly, L. E., & Öz, G. (2023). Antisense Oligonucleotide Silencing Reverses Abnormal Neurochemistry in Spinocerebellar Ataxia 3 Mice. Annals of Neurology, 94(4), 658–671. 10.1002/ana.26713

McLoughlin, H. S., Moore, L. R., Chopra, R., Komlo, R., McKenzie, M., Blumenstein, K. G., Zhao, H., Kordasiewicz, H. B., Shakkottai, V. G., & Paulson, H. L. (2018). Oligonucleotide therapy mitigates disease in spinocerebellar ataxia type 3 mice. Annals of Neurology, 84(1), 64–77. 10.1002/ana.25264

Mendell, J. R., Al-Zaidy, S. A., Lehman, K. J., McColly, M., Lowes, L. P., Alfano, L. N., Reash, N. F., Iammarino, M. A., Church, K. R., Kleyn, A., Meriggioli, M. N., & Shell, R. (2021). Five-Year Extension Results of the Phase 1 START Trial of Onasemnogene Abeparvovec in Spinal Muscular Atrophy. JAMA Neurology, 78(7), 834–841. 10.1001/jamaneurol.2021.1272

Mendell, J. R., Sahenk, Z., Lehman, K., Nease, C., Lowes, L. P., Miller, N. F., Iammarino, M. A., Alfano, L. N., Nicholl, A., Al-Zaidy, S., Lewis, S., Church, K., Shell, R., Cripe, L. H., Potter, R. A., Griffin, D. A., Pozsgai, E., Dugar, A., Hogan, M., & Rodino-Klapac, L. R. (2020). Assessment of Systemic Delivery of rAAVrh74.MHCK7.micro-dystrophin in Children With Duchenne Muscular Dystrophy: A Nonrandomized Controlled Trial. JAMA Neurology, 77(9), 1122–1131. 10.1001/jamaneurol.2020.1484

Mendonça, L. S., Nóbrega, C., Tavino, S., Brinkhaus, M., Matos, C., Tomé, S., Moreira, R., Henriques, D., Kaspar, B. K., & Pereira De Almeida, L. (2019). Ibuprofen enhances synaptic function and neural progenitors proliferation markers and improves neuropathology and motor coordination in Machado-Joseph disease models. Human Molecular Genetics, 28(22), 3691–3703. 10.1093/hmg/ddz097

Miller, V. M., Xia, H., Marrs, G. L., Gouvion, C. M., Lee, G., Davidson, B. L., & Paulson, H. L. (2003). Allele-specific silencing of dominant disease genes. Proceedings of the National Academy of Sciences, 100(12), 7195–7200. 10.1073/pnas.1231012100

Moore, L. R., Rajpal, G., Dillingham, I. T., Qutob, M., Blumenstein, K. G., Gattis, D., Hung, G., Kordasiewicz, H. B., Paulson, H. L., & McLoughlin, H. S. (2017). Evaluation of Antisense Oligonucleotides Targeting ATXN3 in SCA3 Mouse Models. Molecular Therapy - Nucleic Acids, 7(June), 200–210. 10.1016/j.omtn.2017.04.005

Nascimento-Ferreira, I., Nóbrega, C., Vasconcelos-Ferreira, A., Onofre, I., Albuquerque, D., Aveleira, C., Hirai, H., Nicole, D., & Pereira de Almeida, L. (2013). Beclin 1 mitigates motor and neuropathological deficits in genetic mouse models of Machado-Joseph disease. Brain: A Journal of Neurology, 136, 2173–2188. 10.1093/brain/awt144

Nascimento-Ferreira, I., Santos-Ferreira, T., Sousa-Ferreira, L., Auregan, G., Onofre, I., Alves, S., Dufour, N., Gould, V., Koeppen, A., Nicole, D., & Pereira de Almeida, L. (2011). Overexpression of the autophagic beclin-1 protein clears mutant ataxin-3 and alleviates Machado-Joseph disease. Brain: A Journal of Neurology, 134, 1400–1415. 10.1093/brain/awr047

Nobre, R. J., Lobo, D. D., Henriques, C., Duarte, S. P., Lopes, S. M., Silva, A. C., Lopes, M. M., Mariet, F., Schwarz, L. K., Baatje, M. S., Ferreira, V., Vallès, A., Pereira De Almeida, L., Evers, M. M., & Toonen, L. J. A. (2022). MiRNA-Mediated Knockdown of ATXN3 Alleviates Molecular Disease Hallmarks in a Mouse Model for Spinocerebellar Ataxia Type 3. Nucleic Acid Therapeutics, 32(3), 194–205. 10.1089/nat.2021.0020

Nobre, R. J., & Pereira De Almeida, L. (2020). Double stranded rna and uses thereof (World Intellectual Property Association. Patent WO/2020/144611).

Nóbrega, C., Carmo-Silva, S., Albuquerque, D., Vasconcelos-Ferreira, A., Vijayakumar, U.-G., Mendonça, L., Hirai, H., & Pereira de Almeida, L. (2015). Re-establishing ataxin-2 downregulates translation of mutant ataxin-3 and alleviates Machado–Joseph disease. Brain, 138(12), 3537–3554. 10.1093/brain/awv298

Nóbrega, C., Codêsso, J. M., Mendonça, L., & Pereira de Almeida, L. (2019). RNA Interference Therapy for Machado-Joseph Disease: Long-Term Safety Profile of Lentiviral Vectors Encoding Short Hairpin RNAs Targeting Mutant Ataxin-3. Human Gene Therapy, 30(7), 841–854. 10.1089/hum.2018.157

Nóbrega, C., Mendonça, L., Marcelo, A., Lamazière, A., Tomé, S., Despres, G., Matos, C. A., Mechmet, F., Langui, D., den Dunnen, W., de Almeida, L. P., Cartier, N., & Alves, S. (2019). Restoring brain cholesterol turnover improves autophagy and has therapeutic potential in mouse models of spinocerebellar ataxia. Acta Neuropathologica, 138(5), 837–858. 10.1007/s00401-019-02019-7

Nóbrega, C., Nascimento-Ferreira, I., Onofre, I., Albuquerque, D., Déglon, N., & de Almeida, L. P. (2014). RNA interference mitigates motor and neuropathological deficits in a cerebellar mouse model of Machado-Joseph disease. PloS One, 9(8), e100086. 10.1371/journal.pone.0100086

Nóbrega, C., Nascimento-Ferreira, I., Onofre, I., Albuquerque, D., Hirai, H., Déglon, N., & de Almeida, L. P. (2013). Silencing Mutant Ataxin-3 Rescues Motor Deficits and Neuropathology in Machado-Joseph Disease Transgenic Mice. PLoS ONE, 8(1), e52396. 10.1371/journal.pone.0052396

Onofre, I., Mendonça, N., Lopes, S., Nobre, R., De Melo, J. B., Carreira, I. M., Januário, C., Gonçalves, A. F., & De Almeida, L. P. (2016). Fibroblasts of Machado Joseph Disease patients reveal autophagy impairment. Scientific Reports, 6(December 2015), 1–10. 10.1038/srep28220

Orr, H. T., & Zoghbi, H. Y. (2007). Trinucleotide repeat disorders. Annual Review of Neuroscience, 30, 575–621. 10.1146/annurev.neuro.29.051605.113042

Palfi, S., Brouillet, E., Jarraya, B., Bloch, J., Jan, C., Shin, M., Condé, F., Li, X. J., Aebischer, P., Hantraye, P., & Déglon, N. (2007). Expression of mutated huntingtin fragment in the putamen is sufficient to produce abnormal movement in non-human primates. Molecular Therapy, 15(8), 1444–1451. 10.1038/sj.mt.6300185

Paulson, H. L., Das, S. S., Crino, P. B., Perez, M. K., Patel, S. C., Gotsdiner, D., Fischbeck, K. H., & Pittman, R. N. (1997). Machado-Joseph disease gene product is a cytoplasmic protein widely expressed in brain. Annals of Neurology, 41(4), 453–462. 10.1002/ana.410410408

Paulson, H., & Shakkottai, V. (1993). Spinocerebellar Ataxia Type 3. In M. P. Adam, J. Feldman, G. M. Mirzaa, R. A. Pagon, S. E. Wallace, L. J. Bean, K. W. Gripp, & A. Amemiya (Eds.), GeneReviews®. University of Washington, Seattle. http://www.ncbi.nlm.nih.gov/books/NBK1196/

Perelman, P., Johnson, W. E., Roos, C., Seuánez, H. N., Horvath, J. E., Moreira, M. A. M., Kessing, B., Pontius, J., Roelke, M., Rumpler, Y., Schneider, M. P. C., Silva, A., O’Brien, S. J., & Pecon-Slattery, J. (2011). A molecular phylogeny of living primates. PLoS Genetics, 7(3), 1–17. 10.1371/journal.pgen.1001342

Perez, B. A., Shutterly, A., Chan, Y. K., Byrne, B. J., & Corti, M. (2020). Management of neuroinflammatory responses to AAV-mediated gene therapies for neurodegenerative diseases. Brain Sciences, 10(2). 10.3390/brainsci10020119

Prasad, S., Dimmock, D. P., Greenberg, B., Walia, J. S., Sadhu, C., Tavakkoli, F., & Lipshutz, G. S. (2022). Immune Responses and Immunosuppressive Strategies for Adeno-Associated Virus-Based Gene Therapy for Treatment of Central Nervous System Disorders: Current Knowledge and Approaches. Human Gene Therapy, 33(23–24), 1228–1245. 10.1089/hum.2022.138

Rohlfing, T., Kroenke, C. D., Sullivan, E. V., Dubach, M. F., Bowden, D. M., Grant, K., & Pfefferbaum, A. (2012). The INIA19 Template and NeuroMaps Atlas for Primate Brain Image Parcellation and Spatial Normalization. Frontiers in Neuroinformatics, 6. 10.3389/fninf.2012.00027

Rosenberg, R. N. (1992). Machado-Joseph disease: An autosomal dominant motor system degeneration. Movement Disorders, 7(3), 193–203. 10.1002/mds.870070302

Rufino-Ramos, D., R. Albuquerque, P., Leandro, K., Carmona, V., Martins, I., Fernandes, R., Henriques, C., Lobo, D., Faro, R., Perfeito, R., Mendonça, L., Pereira, D., Gomes, C., Nobre, R., & Pereira de Almeida, L. (2023). Extracellular vesicle-based delivery of silencing sequences for the treatment of Machado-Joseph disease/ Spinocerebellar Ataxia type 3. Molecular Therapy: The Journal of the American Society of Gene Therapy, 31. 10.1016/j.ymthe.2023.04.001

Samaranch, L., Salegio, E. A., San Sebastian, W., Kells, A. P., Bringas, J. R., Forsayeth, J., & Bankiewicz, K. S. (2013). Strong cortical and spinal cord transduction after AAV7 and AAV9 delivery into the cerebrospinal fluid of nonhuman primates. Human Gene Therapy, 24(5), 526–532. 10.1089/hum.2013.005

Santana, M. M., Paixão, S., Cunha-Santos, J., Silva, T. P., Trevino-Garcia, A., Gaspar, L. S., Nóbrega, C., Nobre, R. J., Cavadas, C., Greif, H., & Pereira De Almeida, L. (2020). Trehalose alleviates the phenotype of Machado-Joseph disease mouse models. Journal of Translational Medicine, 18(1), 161. 10.1186/s12967-020-02302-2

Saraiva, J., Nobre, R. J., & Pereira de Almeida, L. (2016). Gene therapy for the CNS using AAVs: The impact of systemic delivery by AAV9. Journal of Controlled Release, 241, 94–109. 10.1016/j.jconrel.2016.09.011

Scheel, H. (2003). Elucidation of ataxin-3 and ataxin-7 function by integrative bioinformatics. Human Molecular Genetics, 12(21), 2845–2852. 10.1093/hmg/ddg297

Schmidt, J., & Schmidt, T. (2018). Animal Models of Machado-Joseph Disease. In C. Nóbrega & L. Pereira de Almeida (Eds.), Polyglutamine Disorders (pp. 289–308). Springer International Publishing. 10.1007/978-3-319-71779-1_15

Schmitt, I., Linden, M., Khazneh, H., Evert, B. O., Breuer, P., Klockgether, T., & Wuellner, U. (2007). Inactivation of the mouse Atxn3 (ataxin-3) gene increases protein ubiquitination. Biochemical and Biophysical Research Communications, 362(3), 734– 739. 10.1016/j.bbrc.2007.08.062

Schöls, L., Bauer, P., Schmidt, T., Schulte, T., & Riess, O. (2004). Autosomal dominant cerebellar ataxias: Clinical features, genetics, and pathogenesis. The Lancet. Neurology, 3(5), 291–304. 10.1016/S1474-4422(04)00737-9

Schuster, K. H., Zalon, A. J., DiFranco, D. M., Putka, A. F., Stec, N. R., Jarrah, S. I., Naeem, A., Haque, Z., Zhang, H., Guan, Y., & McLoughlin, H. S. (2024). ASOs are an effective treatment for disease-associated oligodendrocyte signatures in premanifest and symptomatic SCA3 mice. Molecular Therapy, 32(5), 1359–1372. 10.1016/j.ymthe.2024.02.033

Schuster, K. H., Zalon, A. J., Zhang, H., DiFranco, D. M., Stec, N. R., Haque, Z., Blumenstein, K. G., Pierce, A. M., Guan, Y., Paulson, H. L., & McLoughlin, H. S. (2022). Impaired Oligodendrocyte Maturation Is an Early Feature in SCA3 Disease Pathogenesis. The Journal of Neuroscience, 42(8), 1604–1617. 10.1523/JNEUROSCI.1954-20.2021

Simões, A. T., Carmona, V., Duarte-Neves, J., Cunha-Santos, J., & Pereira de Almeida, L. (2022). Identification of the calpain-generated toxic fragment of ataxin-3 protein provides new avenues for therapy of Machado-Joseph disease| Spinocerebellar ataxia type 3. Neuropathology and Applied Neurobiology, 48(1), e12748. 10.1111/nan.12748

Simões, A. T., Gonçalves, N., Koeppen, A., Déglon, N., Kügler, S., Duarte, C. B., & Pereira De Almeida, L. (2012). Calpastatin-mediated inhibition of calpains in the mouse brain prevents mutant ataxin 3 proteolysis, nuclear localization and aggregation, relieving Machado-Joseph disease. Brain, 135(8), 2428–2439. 10.1093/brain/aws177

Simões, A. T., Gonçalves, N., Nobre, R. J., Duarte, C. B., & Pereira de Almeida, L. (2014). Calpain inhibition reduces ataxin-3 cleavage alleviating neuropathology and motor impairments in mouse models of machado-Joseph disease. Human Molecular Genetics, 23(18), 4932–4944. 10.1093/hmg/ddu209

Song, G., Ma, Y., Gao, X., Zhang, X., Zhang, F., Tian, C., Hou, J., Liu, Z., Zhao, Z., & Tian, Y. (2022). CRISPR/Cas9-mediated genetic correction reverses spinocerebellar ataxia 3 disease-associated phenotypes in differentiated cerebellar neurons. Life Medicine, 1(1), 27–44. 10.1093/lifemedi/lnac020

Switonski, P. M., Fiszer, A., Kazmierska, K., Kurpisz, M., Krzyzosiak, W. J., & Figiel, M. (2011). Mouse Ataxin-3 Functional Knock-Out Model. Neuromolecular Medicine, 13(1), 54–65. 10.1007/s12017-010-8137-3

Tai, C.-H., Lee, N.-C., Chien, Y.-H., Byrne, B. J., Muramatsu, S.-I., Tseng, S.-H., & Hwu, W.-L. (2022). Long-term efficacy and safety of eladocagene exuparvovec in patients with AADC deficiency. Molecular Therapy: The Journal of the American Society of Gene Therapy, 30(2), 509–518. 10.1016/j.ymthe.2021.11.005

Takiyama, Y., Nishizawa, M., Tanaka, H., Kawashima, S., Sakamoto, H., Karube, Y., Shimazaki, H., Soutome, M., Endo, K., Ohta, S., Kagawa, Y., Kanazawa, I., Mizuno, Y., Yoshida, M., Yuasa, T., Horikawa, Y., Oyanagi, K., Nagai, H., Kondo, T.,… Tsuji, S. (1993). The gene for Machado-Joseph disease maps to human chromosome 14q. Nature Genetics, 4(3), 300–304. 10.1038/ng0793-300

Tarantal, A. F., Noctor, S. C., & Hartigan-O’Connor, D. J. (2022). Nonhuman Primates in Translational Research. Annual Review of Animal Biosciences, 10, 441–468. 10.1146/annurev-animal-021419-083813

Tomioka, I., Ishibashi, H., Minakawa, E. N., Motohashi, H. H., Takayama, O., Saito, Y., Popiel, H. A., Puentes, S., Owari, K., Nakatani, T., Nogami, N., Yamamoto, K., Noguchi, S., Yonekawa, T., Tanaka, Y., Fujita, N., Suzuki, H., Kikuchi, H., Aizawa, S.,… Seki, K. (2017). Transgenic monkey model of the polyglutamine diseases recapitulating progressive neurological symptoms. eNeuro, 4(2), 1–16. 10.1523/ENEURO.0250-16.2017

Tomioka, I., Nogami, N., Nakatani, T., Owari, K., Fujita, N., Motohashi, H., Takayama, O., Takae, K., Nagai, Y., & Seki, K. (2017). Generation of transgenic marmosets using a tetracyclin-inducible transgene expression system as a neurodegenerative disease model. Biology of Reproduction, 97(5), 772–780. 10.1093/biolre/iox129

Toonen, L., Schmidt, I., Luijsterburg, M., van Attikum, H., & van Roon-Mom, W. (2016). Antisense oligonucleotide-mediated exon skipping as a strategy to reduce proteolytic cleavage of ataxin-3. Scientific Reports, 6, 35200. 10.1038/srep35200

Vallender, E. J., & Miller, G. M. (2013). Nonhuman Primate Models in the Genomic Era: A Paradigm Shift. ILAR Journal, 54(2), 154–165. 10.1093/ilar/ilt044

Vasconcelos-Ferreira, A., Martins, I. M., Lobo, D., Pereira, D., Lopes, M. M., Faro, R., Lopes, S. M., Verbeek, D., Schmidt, T., Nóbrega, C., & Pereira de Almeida, L. (2022). ULK overexpression mitigates motor deficits and neuropathology in mouse models of Machado-Joseph disease. Molecular Therapy: The Journal of the American Society of Gene Therapy, 30(1), 370–387. 10.1016/j.ymthe.2021.07.012

Warrick, J. M., Paulson, H. L., Gray-Board, G. L., Bui, Q. T., Fischbeck, K. H., Pittman, R. N., & Bonini, N. M. (1998). Expanded polyglutamine protein forms nuclear inclusions and causes neural degeneration in Drosophila. Cell, 93(6), 939–949. 10.1016/s0092-8674(00)81200-3

Watchon, M., Yuan, K. C., Mackovski, N., Svahn, A. J., Cole, N. J., Goldsbury, C., Rinkwitz, S., Becker, T. S., Nicholson, G. A., & Laird, A. S. (2017). Calpain Inhibition Is Protective in Machado-Joseph Disease Zebrafish Due to Induction of Autophagy. The Journal of Neuroscience: The Official Journal of the Society for Neuroscience, 37(32), 7782– 7794. 10.1523/JNEUROSCI.1142-17.2017

Weishäupl, D., Schneider, J., Peixoto Pinheiro, B., Ruess, C., Dold, S. M., von Zweydorf, F., Gloeckner, C. J., Schmidt, J., Riess, O., & Schmidt, T. (2019). Physiological and pathophysiological characteristics of ataxin-3 isoforms. Journal of Biological Chemistry, 294(2), 644–661. 10.1074/jbc.RA118.005801

Weiss, A. R., Liguore, W. A., Brandon, K., Wang, X., Liu, Z., Domire, J. S., Button, D., Srinivasan, S., Kroenke, C. D., & McBride, J. L. (2022). A novel rhesus macaque model of Huntington’s disease recapitulates key neuropathological changes along with motor and cognitive decline. eLife, 11, e77568. 10.7554/eLife.77568

Wetzel, J. S., Hardcastle, N., Tora, M. S., Federici, T., Frey, S., Novek, J., Arulanandam, T., Johnson, M., Pielemeier, R., & Boulis, N. M. (2019). Frameless Stereotactic Targeting of the Cerebellar Dentate Nucleus in Nonhuman Primates: Translatable Model for the Surgical Delivery of Gene Therapy. Stereotactic and Functional Neurosurgery, 97(5– 6), 293–302. 10.1159/000504858

Wiseman, J. P., Scarrott, J. M., Alves-Cruzeiro, J., Saffari, A., Böger, C., Karyka, E., Dawes, E., Davies, A. K., Marchi, P. M., Graves, E., Fernandes, F., Yang, Z.-L., Coldicott, I., Hirst, J., Webster, C. P., Highley, J. R., Hackett, N., Angyal, A., Silva, T. de,… Azzouz, M. (2024). Pre-clinical development of AP4B1 gene replacement therapy for hereditary spastic paraplegia type 47. EMBO Molecular Medicine. 10.1038/s44321-024-00148-5

Yang, W., Wang, G., Wang, C. E., Guo, X., Yin, P., Gao, J., Tu, Z., Wang, Z., Wu, J., Hu, X., Li, S., & Li, X. J. (2015). Mutant alpha-synuclein causes age-dependent neuropathology in monkey brain. Journal of Neuroscience, 35(21), 8345–8358. 10.1523/JNEUROSCI.0772-15.2015

Yin, P., Guo, X., Yang, W., Yan, S., Yang, S., Zhao, T., Sun, Q., Liu, Y., Li, S., & Li, X.-J. (2019). Caspase-4 mediates cytoplasmic accumulation of TDP-43 in the primate brains. Acta Neuropathologica, 137(6), 919–937. 10.1007/s00401-019-01979-0

