## Supplemental material for "AAV-mediated allele-specific silencing alleviates neuropathology in a novel non-human primate model of Spinocerebellar ataxia type 3"

**Author names and affiliations:**

Carina Henriques[1-5], Diana Lobo[1-3,5-6], Ana Carolina Silva[1-3,5-6], Ana Rita Fernandes[1-2,5-6], Miguel Monteiro Lopes[1-3,5-6], Audrey Fayard[7-8], Caroline Jan[7-8], Sophie Lecourtois[7-8], Martine Guillermier[7-8], Julien Flament[7-8], João Castelhano[9-11], Miguel Castelo-Branco[9-11], Rui Caetano Oliveira[12-13], Philippe Hantraye[7-8], Padmaja Yalamanchili[14], Steven de Marco[14], Romina Aron Badin[7-8,†], Luís Pereira de Almeida[1-5,†], Rui Jorge Nobre[1-3,5-6,†]

1 Center for Neuroscience and Cell Biology (CNC), Gene and Stem Cell Therapies for the Brain Group, University of Coimbra, 3004-504 Coimbra, Portugal

2 Center for Innovative Biomedicine and Biotechnology (CIBB), Vectors, Gene and Cell Therapy Group, University of Coimbra, 3004-504 Coimbra, Portugal

3 ViraVector–Viral Vector for Gene Transfer Core Facility, University of Coimbra, 3004-504 Coimbra, Portugal

4 Faculty of Pharmacy, University of Coimbra, 3000-548 Coimbra, Portugal

5 GeneT, Center for Excellence in Gene Therapy in Portugal, University of Coimbra, 3004-504 Coimbra, Portugal

6 Institute for Interdisciplinary Research (III), University of Coimbra, 3030-789 Coimbra, Portugal

7 CEA, DRF, Institute of Biology François Jacob, Molecular Imaging Research Center (MIRCen), 92265 Fontenay-aux-Roses, France

8 CNRS, CEA, Paris-Sud University, Université Paris-Saclay, Neurodegenerative Diseases Laboratory (UMR9199), 92265 Fontenay-aux-Roses, France

9 Coimbra Institute for Biomedical Imaging and Translational Research (CIBIT), University of Coimbra, 3000-548 Coimbra, Portugal

10 Institute for Nuclear Sciences Applied to Health (ICNAS), University of Coimbra, 3000-548 Coimbra, Portugal

11 Faculty of Medicine, University of Coimbra, 3000-370 Coimbra, Portugal

12 Germano de Sousa Pathological Anatomy Center, 3000-377, Coimbra, Portugal

13 Institute of Histology and Embryology, Faculty of Medicine, University of Coimbra, 3004-504, Coimbra, Portugal

14 PTC Therapeutics, 500 Warren Corp Center Dr, Warren, NJ 07059, USA

†These are corresponding authors and contributed equally to this work as senior authors.

**Corresponding Authors:**

Correspondence to: Luís Pereira de Almeida and Rui Jorge Nobre

CNC - Center for Neuroscience and Cell Biology, University of Coimbra, Rua Larga, 3004-504 Coimbra, Portugal

### Supplemental Material

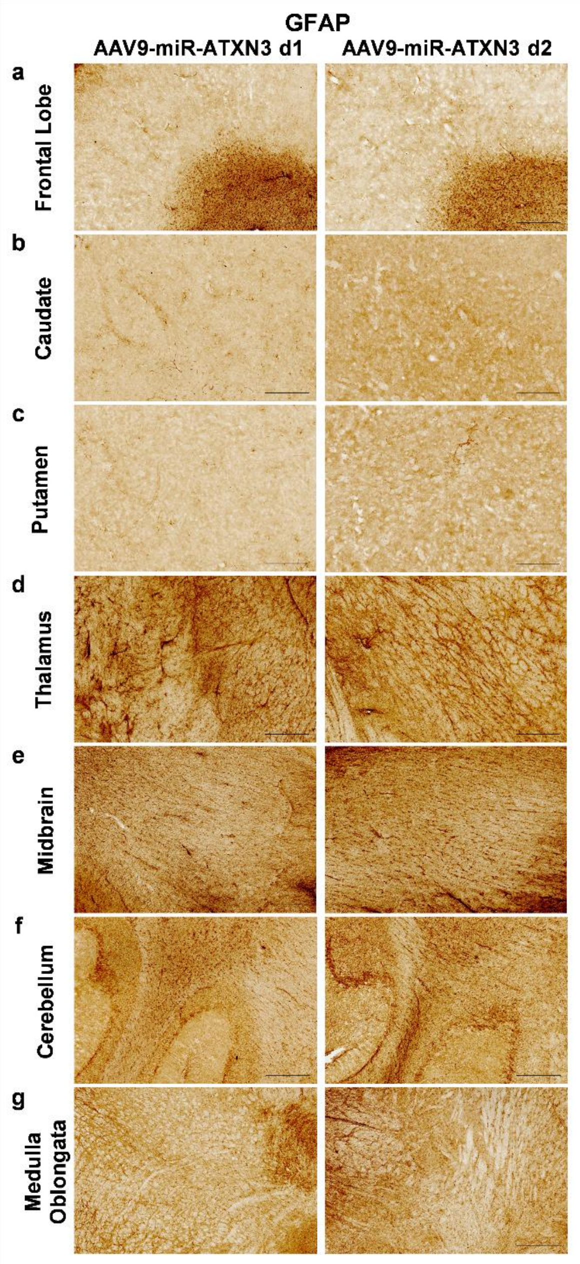


**Supplementary Figure 1 - No major astrogliosis was observed in the NHP brains following ICM delivery for either dose of AAV9-miR-ATXN3.**

Immunohistochemistry for GFAP in the frontal lobe **(a)**, caudate **(b)**, putamen **(c)**, thalamus **(d)**, midbrain **(e)**, cerebellum **(f)**, medulla oblongata **(g)** of monkeys injected with AAV9-miR-ATXN3 d1 and AAV9-miR-ATXN3. **(a-g)** Monkeys' coronal brain sections were incubated with GFAP antibody. Scale bar = 500μm.

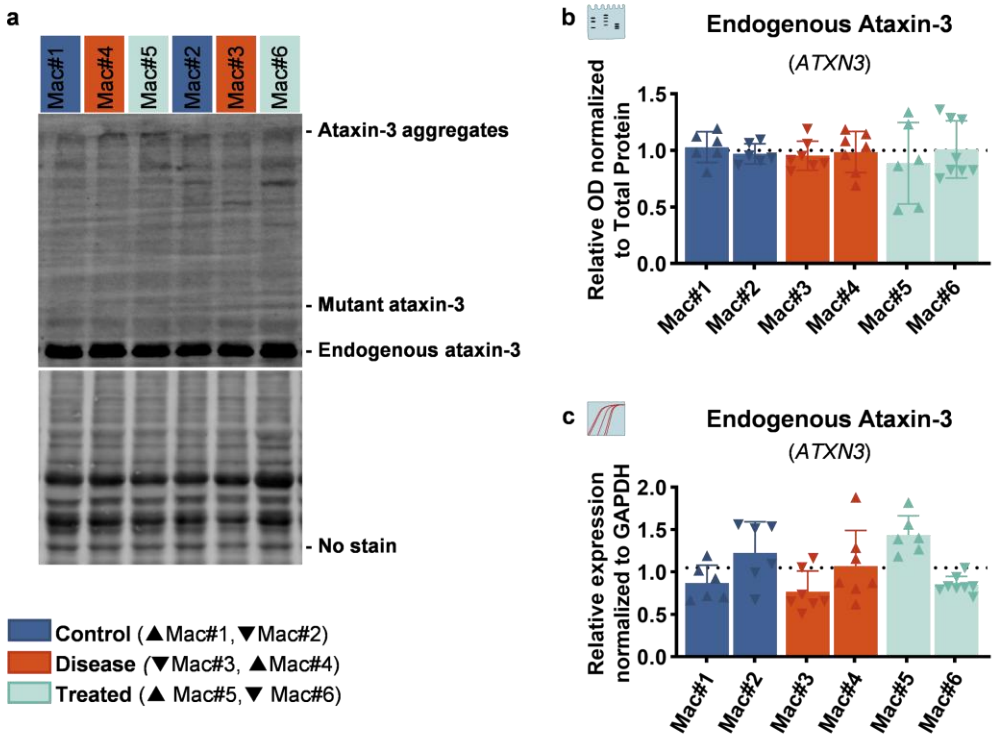


**Supplementary Figure 2 - No differences were observed in endogenous *Macaca fascicularis* Ataxin-3 protein and mRNA levels.**

**(a)** Membranes of cerebellum extracts were incubated with Ataxin-3 antibody 1h9 and total protein (No stain) was used for densitometric normalisation. (**b**) Densitometric quantification of *Macaca fascicularis* endogenous Ataxin-3 protein in the cerebellum relative to total protein densitometry. **(c)** Expression levels of endogenous Ataxin-3 (*Macaca fascicularis*) (*ATXN3*) mRNA, relative to *Macaca fascicularis GAPDH.* Results are shown as mean ± SD for **(b,c)** control, mut*ATXN3*, and mut*ATXN3* + AAV9-miR-ATXN3 (n = 6-8, per animal). The dotted line represents the average of the control group.

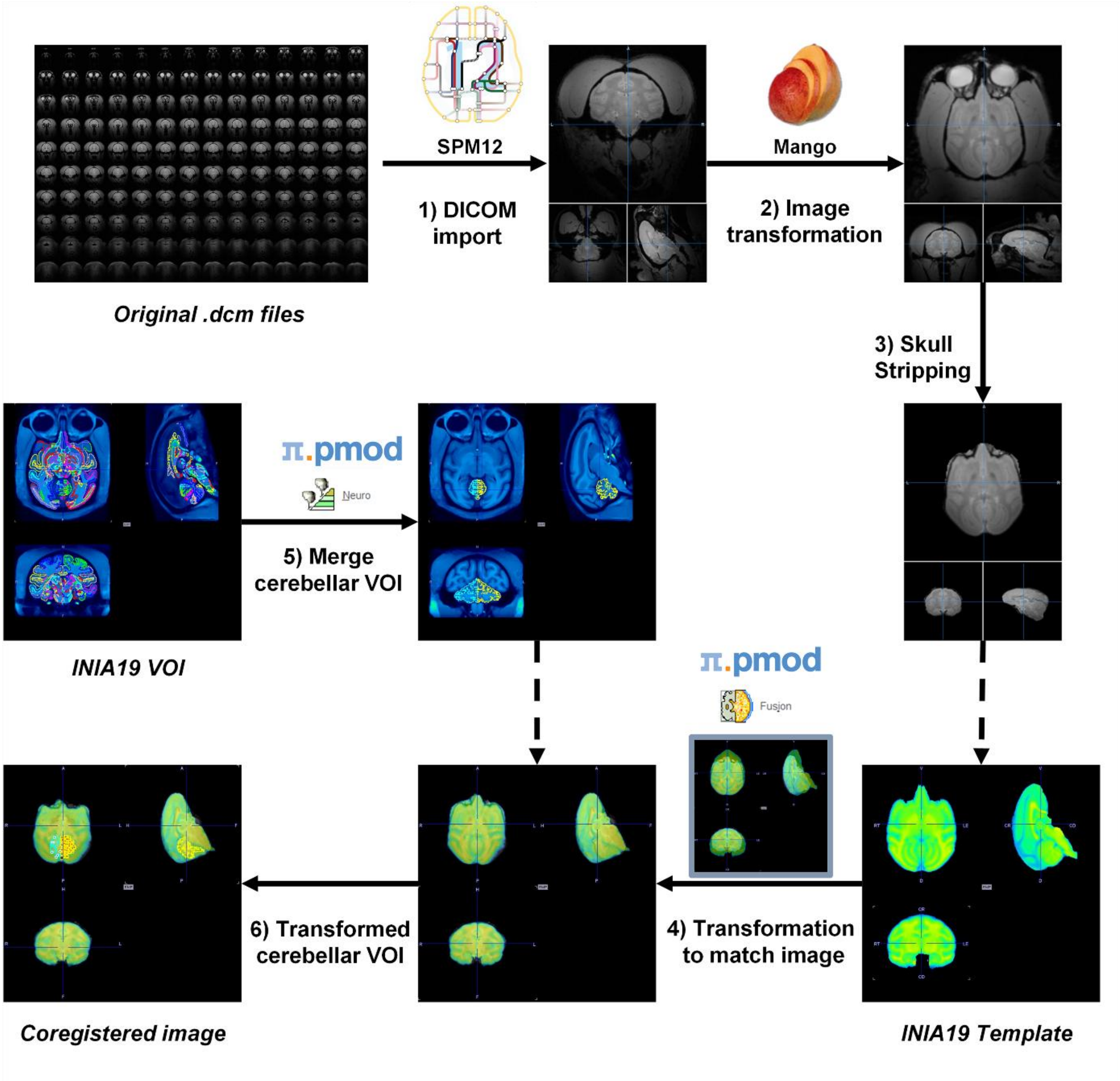


**Supplementary Figure 3 - Overview of the pipeline for MRI analysis.**
Firstly, (1) T2 DCIOM images were converted to NIFTI-1 using the SPM12 DICOM import function. Then, using Mango software, (2) images were transformed to the desired orientation and (3) the skull was stripped. Using PMOD module Fusion, (4) the INIA19 template was transformed to match our skull-stripped images. Finally, the transformation applied to the INIA19 template was applied to the cerebellar VOI, previously (5) created by the fusion of all INIA19 cerebellar VOI in the PMOD module Neuro. Cerebellar volumes were obtained by (6) the coregistration of the transformed cerebellar VOI with the transformed INIA19 template.

**Supplementary Table 1 - Experimental design for AAV9-miR-ATXN3 biodistribution experiment.**

| Experimental Group | N of animals | RoA | Agent | Dose/animal | Injection volume/animal |
|---|---|---|---|---|---|
| **AAV9-miR-ATXN3 (d1)** | 3 | ICM | scAAV9-miR-ATXN3 | $0.3 \times 10^{13}$ vg | 1.5 mL |
| **AAV9-miR-ATXN3 (d2)** | 3 | ICM | scAAV9-miR-ATXN3 | $3 \times 10^{13}$ vg | 1.5 mL |

ICM = intracisterna magna; N =number; scAAV9-miR-ATXN3 (abbreviation: AAV9-miR-ATXN3) = adeno-associated viral vectors, serotype 9, with a self-complementary (sc) genome, encoding for microRNA (miRNA) for the silencing of mutant Ataxin-3 (miR-ATXN3); RoA =route of administration; vg = viral genomes.

**Supplementary Table 2 - Experimental design for AAV9-miR-ATXN3 target engagement experiment.**

| Experimental Group | N of animals | RoA | Agent | Dose/animal | Injection volume/animal |
|---|---|---|---|---|---|
| **Control group** | 2 | IP – WM, near DCNs | PBS/BSA 1% | - | 6 IS x 15 µL |
| | | ICM | PBS/0.001% F68 | - | 1.5 mL |
| **Disease group** | 2 | IP – WM, near DCNs | LV-mut*ATXN3*-Q72 | 20 000 ng of P24 | 6 IS x 15 µL |
| | | ICM | PBS/0.001% F68 | - | 1.5 mL |
| **Treated group** | 2 | IP – WM, near DCNs | LV-mut*ATXN3*-Q72 | 20 000 ng of P24 | 6 IS x 15 µL - |
| | | ICM | scAAV9-miR-ATXN3 | 2 x $10^{13}$ vg | 1.5 mL |

BSA = bovine serum albumin; DCN = deep cerebellar nuclei; F68 = pluronic F-68; ICM = intracisterna magna; IP = intraparenchymal; IS = injection site; LV-mut*ATXN3* = lentiviral vectors encoding for mutant Ataxin-3; N =number; PBS = phosphate-buffered saline; scAAV9-miR-ATXN3 (abbreviation: AAV9-miR-ATXN3) = adeno-associated viral vectors, serotype 9, with a self-complementary (sc) genome, encoding for microRNA (miRNA) for the silencing of mutant Ataxin-3 (miR-ATXN3); RoA =route of administration; vg = viral genomes; WM = white matter.

#### Supplementary Table 3 - Primers used in real-time PCR.

| | | Real-time PCR Primers | |
|---|---|---|---|
| **Gene** | **Specie** | **Sequence** (5'→3') | **Ta** (°C) |
| ***ATXN3-Q72*** | *Homo sapiens* | **F:** AGTCCATCTTCCACGAGAAACAAGA<br>**R:** TCCACAGGGCTAAAATATTCTCCT | 58 |
| ***ATXN3*** | *Macaca fascicularis* | **F:** ATTCAGCTAAGTATGCAAGGTAGT<br>**R:** AGATTTGTACCTGATGTCTGTGGA | 56 |
| ***CALB1*** | *Macaca fascicularis* | **F:** CCTGCAATCGTCCCTCATCA<br>**R:** CCAGGTAACCACTTCCGTCA | 56 |
| ***GAPDH*** | *Macaca fascicularis* | **F:** ACAACAGCCTCAAGATCGTCAG<br>**R:** ACTGTGGTCATGAGTCCTTCC | 56 |
| ***GFAP*** | *Macaca fascicularis* | **F:** AGCCTGGACACCAAGTCTGT<br>**R:** GGACTCCTTAATGACCTCTCCA | 54 |
| ***MAPT*** | *Macaca fascicularis* | **F:** AGCCAACGCCACCAGGATT<br>**R:** CTGATTTTGGAGGTTCACCAGAGC | 58 |
| ***MOBP*** | *Macaca fascicularis* | **F:** AAAACGACCAAGGAGGGTCC<br>**R:** ATTGAGGAAGGTGAACGGCG | 56 |
| ***PLP1*** | *Macaca fascicularis* | **F:** TGCAAAACAGCTGAGTTCCAAA<br>**R:** CGGCAAAGTTGTAAGTGGCA | 54 |
| ***RBFOX3*** | *Macaca fascicularis* | **F:** CTACAGCGACAGTTACGGCA<br>**R:** GGTTCCAATGCTGTAGGTTGC | 54 |

F = forward; na = not applicable; R = reverse; Ta = annealing temperature.

**Supplementary Table 4 - Pearson's correlation and simple linear regression analysis between AAV viral genomes and miR-ATXN3 copy number.**

| Experiment and experimental group | Samples | Total number of values (n) | *Pearson's r* | Simple linear regression ($r^2$) | *p value* | Conclusion |
|---|---|---|---|---|---|---|
| **BD:<br>AAV9-miR-ATXN3 d<br>+<br>AAV9-miR-ATXN3 d2** | All dissected regions | 70 | 0.7088 | 0.5024 | < 0.0001 | Strong correlation |
| | Cerebellum | 7 | 0.6659 | 0.4434 | 0.1025 | Tendency for moderate correlation |
| **BD:<br>AAV9-miR-ATXN3 d1** | All dissected regions | 17 | 0.3021 | 0.09124 | 0.2387 | Not enough evidence to conclude |
| | Cerebellum | 2 | - | - | - | - |
| **BD:<br>AAV9-miR-ATXN3 d2** | All dissected regions | 53 | 0.7841 | 0.6148 | <0.0001 | Strong correlation |
| | Cerebellum | 5 | 0.4955 | 0.2455 | 0.3960 | Not enough evidence to conclude |
| **TE:<br>Treated group** | Cerebellum | 11 | 0.1708 | 0.02919 | 0.6155 | Not enough evidence to conclude |

BD = biodistribution; d = dose; TE = target engagement.

#### Supplementary Table 5 - Multivariable correlation matrix between *p* values.

| | Mutant Ataxin-3 | *ATXN3* | *RBFOX3* | *Calb1* | *MAPT* | *GFAP* | *PLP1* | *MOBP* | Cb Volume w12 (% to BL) | Provirus Copies/ μg DNA |
|---|---|---|---|---|---|---|---|---|---|---|
| **Mutant Ataxin-3** | | 0.3616 | 0.1198 | 0.4657 | 0.2110 | 0.1542 | 0.0945 | 0.1215 | 0.1072 | 0.8498 |
| ***ATXN3*** | 0.3616 | | 0.0119 | 0.0673 | 0.0016 | 0.8072 | 0.5652 | 0.3305 | 0.8697 | 0.5591 |
| ***RBFOX3*** | 0.1198 | 0.0119 | | 0.0582 | 0.0016 | 0.7006 | 0.3723 | 0.1718 | 0.5316 | 0.4046 |
| ***Calb1*** | 0.4657 | 0.0673 | 0.0582 | | 0.0450 | 0.5413 | 0.7954 | 0.9311 | 0.6138 | 0.9607 |
| ***MAPT*** | 0.2110 | 0.0016 | 0.0016 | 0.0450 | | 0.7408 | 0.4480 | 0.2625 | 0.7468 | 0.5508 |
| ***GFAP*** | 0.1542 | 0.8072 | 0.7006 | 0.5413 | 0.7408 | | 0.0108 | 0.1014 | 0.0389 | 0.9155 |
| ***PLP1*** | 0.0945 | 0.5652 | 0.3723 | 0.7954 | 0.4480 | 0.0108 | | 0.0085 | 0.0125 | 0.4417 |
| ***MOBP*** | 0.1215 | 0.3305 | 0.1718 | 0.9311 | 0.2625 | 0.1014 | 0.0085 | | 0.0218 | 0.1434 |
| **Cb Volume w12 (% to BL)** | 0.1072 | 0.8697 | 0.5316 | 0.6138 | 0.7468 | 0.0389 | 0.0125 | 0.0218 | | 0.3618 |
| **Provirus Copies/ μg DNA** | 0.8498 | 0.5591 | 0.4046 | 0.9607 | 0.5508 | 0.9155 | 0.4417 | 0.1434 | 0.3618 | |

BL = baseline; Cb = cerebellum; w = week.
